# Molecular principles of CRISPR-Cas13 mismatch intolerance enable selective silencing of point-mutated oncogenic RNA with single-base precision

**DOI:** 10.1101/2023.09.26.557083

**Authors:** Carolyn Shembrey, Ray Yang, Joshua Casan, Wenxin Hu, Teresa Sadras, Krishneel Prasad, Jake Shortt, Ricky W Johnstone, Joseph A Trapani, Paul G Ekert, Mohamed Fareh

## Abstract

Single nucleotide variants (SNVs) are extremely prevalent in human cancers. For instance, KRAS mutations occur in over 90% of pancreatic cancers and ∼40% of colorectal cancers. Virtually all KRAS mutations are SNVs, most of which remain clinically unactionable. The programmable RNA nuclease CRISPR-Cas13 has been deployed to specifically target RNAs such as overexpressed oncogenes and fusion transcripts. However, silencing oncogenic SNVs with single-base precision remains extremely challenging due to the intrinsic mismatch tolerance of Cas13. Here, we developed a comprehensive mutagenesis analysis of target-spacer interactions at single-nucleotide resolution, which revealed key spacer nucleotide positions intolerant to mismatches. We show that introducing synthetic mismatches at these precise positions enables *de nova* design of CRISPR RNA (crRNA) with strong preferential silencing of SNV transcripts. We demonstrate that our top-performing crRNAs possess prominent SNV-selectivity with dose-dependent silencing activity against all KRAS G12 variants at both the RNA and protein levels with minimal off-target silencing of wildtype KRAS. We applied these design principles to effectively silence oncogenic NRAS G12D and BRAF V600E transcripts, underscoring the adaptability of this platform to silence various SNVs. These findings demonstrate that the CRISPR Cas13 system can be reprogrammed to target mutant transcripts with single-base precision, showcasing the tremendous potential of this tool in personalized transcriptome editing.

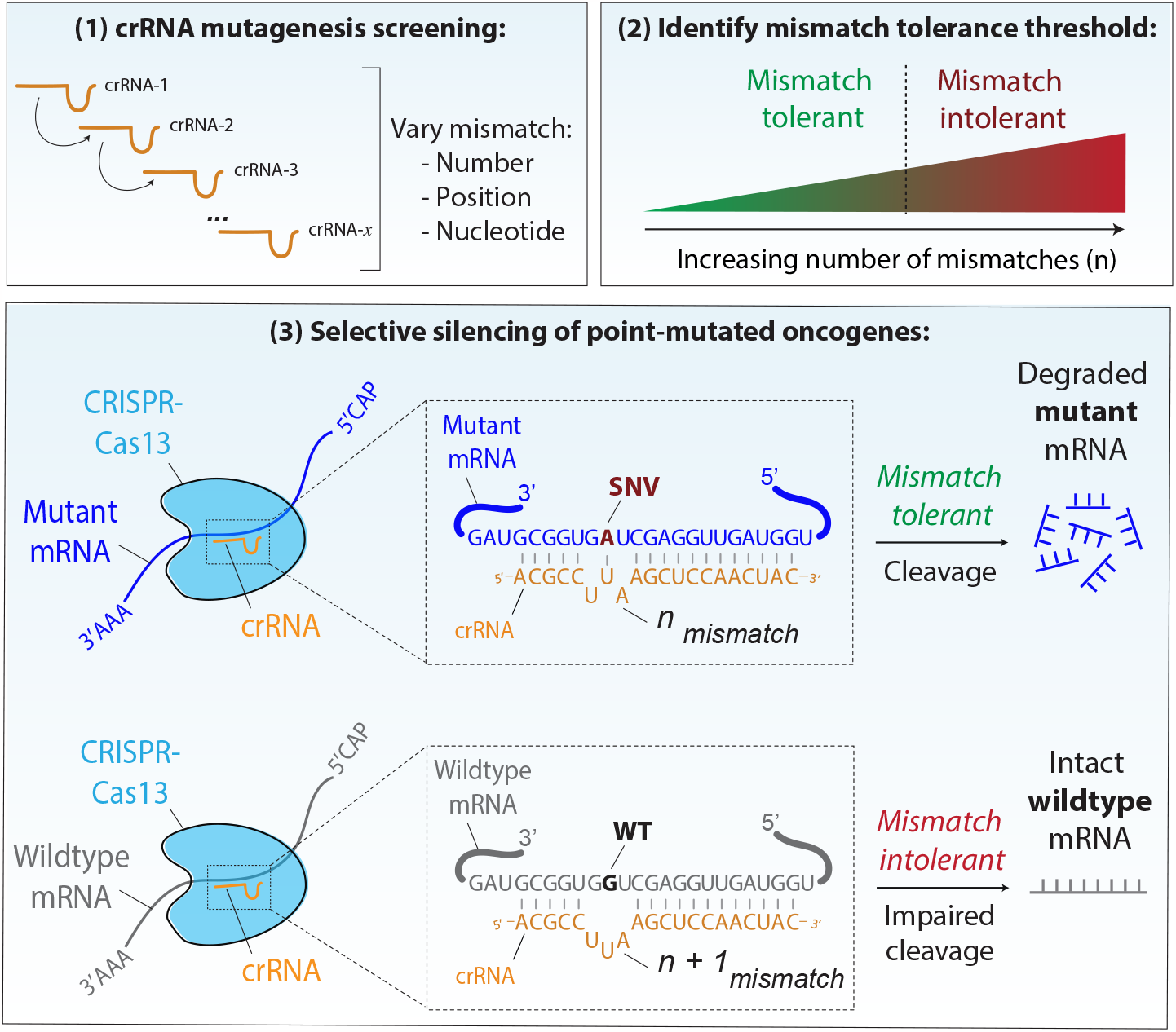

## INTRODUCTION

Comprehensive molecular profiling initiatives have revealed the vast landscape of single-nucleotide variants (SNV) in adult^1^ and paediatric cancers^2^. On average, adult tumours harbour 2.6 driver SNVs^1^ which are often enriched in proliferative and pro-survival pathways such as the mitogen activated protein kinase (MAPK) pathway. Somatic MAPK aberrations most commonly occur in the proto-oncogenes KRAS (Kirsten rat sarcoma viral oncogene homologue) or BRAF (v-raf murine sarcoma viral oncogene homologue Bl), which collectively drive malignancy in approximately one third of all human cancers^1^. KRAS mutations are extremely pervasive, present in over 90% of pancreatic cancers, ∼40% of colorectal cancers and ∼30% of lung cancers^3^. Virtually all (>99%) KRAS mutations are missense substitutions^4^, with the most frequent KRAS SNVs occurring at the codon 12-encoded glycine residue^4^•

Tumour-associated SNVs are considered an attractive therapeutic target due to their high prevalence and restricted expression in transformed cells. Several small molecule inhibitors targeting oncogenic SNVs have been clinically approved, including a KRAS^Giic^ inhibitor (sotorasib). This agent exploits a unique binding pocket generated in the mutated protein, enabling its G12C selectivity^5^’^6^. However, exclusive targeting of other SNVs remains a significant pharmacological challenge as these mutated proteins differ from their wildtype counterparts by only one amino acid, rarely conferring sufficient structural dissimilarity for SNV-selective targeting. Indeed, most KRAS aberrations (such as -G12R, -G13D, - Q61H etc.) lack any accessible binding pockets and are therefore considered clinically unactionable^5^. As only ∼10% of KRAS mutations are of the G12C genotype^7^, therapeutic targeting of non-G12C variants remains an unmet clinical challenge. Although a recent landmark study has developed a pre-clinical, pan-KRAS inhibitor, which suppresses multiple KRAS variants by non selectively trapping the protein in its inactive state ^8^, this new drug also suppresses wildtype KRAS and may therefore exhibit toxicity in normal tissues.

Unfortunately, the reported number of deleterious, tumour-associated SNVs far outweighs the number of clinically approved inhibitors. The development of traditional small molecule inhibitors that target proteins requires structure-based design and high-throughput screens; a process that is expensive, labour-intensive and low-yield. In contrast, new innovative sequence-based therapeutics that target oncogenic drivers at their DNA or RNA level can be developed and personalised for each patient’s specific mutational profile with rapid manufacturing and minimum cost. Additionally, in bypassing any three-dimensional structural constraints, sequence-based therapeutics targeting aberrant nucleic acid molecules might provide a novel modality for targeting SNVs that generate proteins that lack pharmacophoric vulnerabilities.

CRISPR-Cas9 molecular tools have revolutionised our ability to perform targeted genome editing. Elegant studies have exploited the observation that certain SNVs, such as EGFR^LsssR^, generate novel 5’-NGG-3’ protospacer adjacent motif (PAM) sequences recognisable by SpCas9, thus enabling the selective cleavage of the mutated oncogene whilst leaving the PAM-free wildtype sequence intact^9^. However, several reports have linked the on target and off-target nuclease activity of CRISPR-Cas9 to megabase-scale chromosomal truncations, rearrangeme nts, and even chromothripsis at both the target and distal sites^10-13^, as well as significantly increased rates of off target INDEL generation^14^. Cas9 genome editing has also been shown to positively select for cells with defective p53-mediated DNA damage responses and may thus lead to the expansion of TP53 mutated clones from a heterogenous cancer cell population^15^’^16^. Thus, the risks associated with such permanent, off-target genomic alterations may limit the therapeutic use of current CRISPR-Cas9 technologies in clinical settings.

Conversely, CRISPR-Cas13 is an RNA-guided RNA targeting nuclease that enables precise and efficient cleavage of single-stranded RNA without altering genomic DNA^17^. Previous studies have demonstrated that CRISPR-Cas13 can be successfully reprogrammed to silence various pathogenic RNAs including SARS-CoV-2 variants of concern^18,19^ and to alleviate neurodegenerative symptoms in mouse models of Huntington’s disease^20^. Cas13 enzymes typically require basepairing with a 23 to 30-nucleotide (nt) long recognition motif to activate its HEPN domains and cleave the RNA target. This highlights the high specificity of Cas13 enzymes compared to classical eukaryotic RNA interreference (RNAi), where only a 7 to 8-nt long seed region dictates target recognition and degradation^21^. As such, traditional RNAi is highly permissive to off-target binding, with some estimates indicating that over one third of all human transcripts could harbour one or several siRNA binding sites^22^. Conversely, the greater specificity of Cas13 renders it a promising tool for targeting oncogenic transcripts. In fact, we and others have reprogrammed Cas13 to silence oncogenic fusion transcripts by targeting their tumour-specific breakpoint sequence^23,24^. However, single-base precision silencing of SNVs with Cas13 remains highly challenging due to the well-known mismatch tolerance capability of various Cas13 enzymes^18^’^25-27^. Thus, systematic and precise silencing of SNVs may only be achieved once we understand the molecular principles behind Cas13 mismatch tolerance and intolerance at the single-base level.

Here, we present a novel experimental pipeline to design and validate Cas13-compatible CRISPR RNAs (crRNAs) capable of SNV-specific transcript obliteration. By employing an extensive analysis of target-spacer interactions at the level of individual nucleotides, we developed a suite of crRNAs with a remarkable ability to selectively silence point-mutant KRAS transcripts. Our most effective crRNAs demonstrated marked preferential silencing of various oncogenic KRAS mutant transcripts, displaying dose-dependent and SNV-selective silencing. Furthermore, we successfully applied these design principles to silence oncogenic NRAS G12D and BRAF VG00E transcripts, demonstrating the adaptability of this approach for a diverse range of SNV targets. These findings highlight the versatility of our Cas13 design principles to precisely target mutant transcripts with single-base precision.

## RESULTS

### Selective cleavage of KRAS G12C and G12D mutant transcripts via precise spacer mutagenesis

We questioned whether Cas13 can efficiently silence mutant KRAS transcripts while sparing the wildtype variant. Among various Cas13 orthologs, we focused on *Ruminococcus flavefaciens* Cas13d (RfxCas13d) that has been shown to possess high silencing activity ^28^• To assess the RNA cleavage activity and specificity of *RfxCas13d* and its crRNAs against mutants and wildtype KRAS, we utilized a reporter assay that enables accurate monitoring of Cas13-mediated mRNA cleavage via loss-of-fluorescence signal in HEK 293T cells^18^ **(Figure lA, methods)**.

As G12D is the most common KRAS SNV ^4,^ we first designed a parental crRNA (crKRAS) which basepairs with the G12D transcript with 100% sequence complementarity (termed a “perfect-match” crRNA). Here, the spacer’s “U” nucleotide that basepairs with the “A” nucleotide substitution found in G12D-mutated KRAS mRNA is located at crRNA position 10 **(Figure 1B, upper)**. Whilst crKRAS exhibited incredibly potent mRNA cleavage relative to the non-targeting control (crNT), it efficiently silenced both the wildtype KRAS and G12D transcripts with equal efficiency **(Figure 1B, lower)**, indicating that RfxCas13d has an intrinsic single-nucleotide mismatch tolerance that disables the discrimination between WT and SNV tumour transcripts.

**Figure 1.**
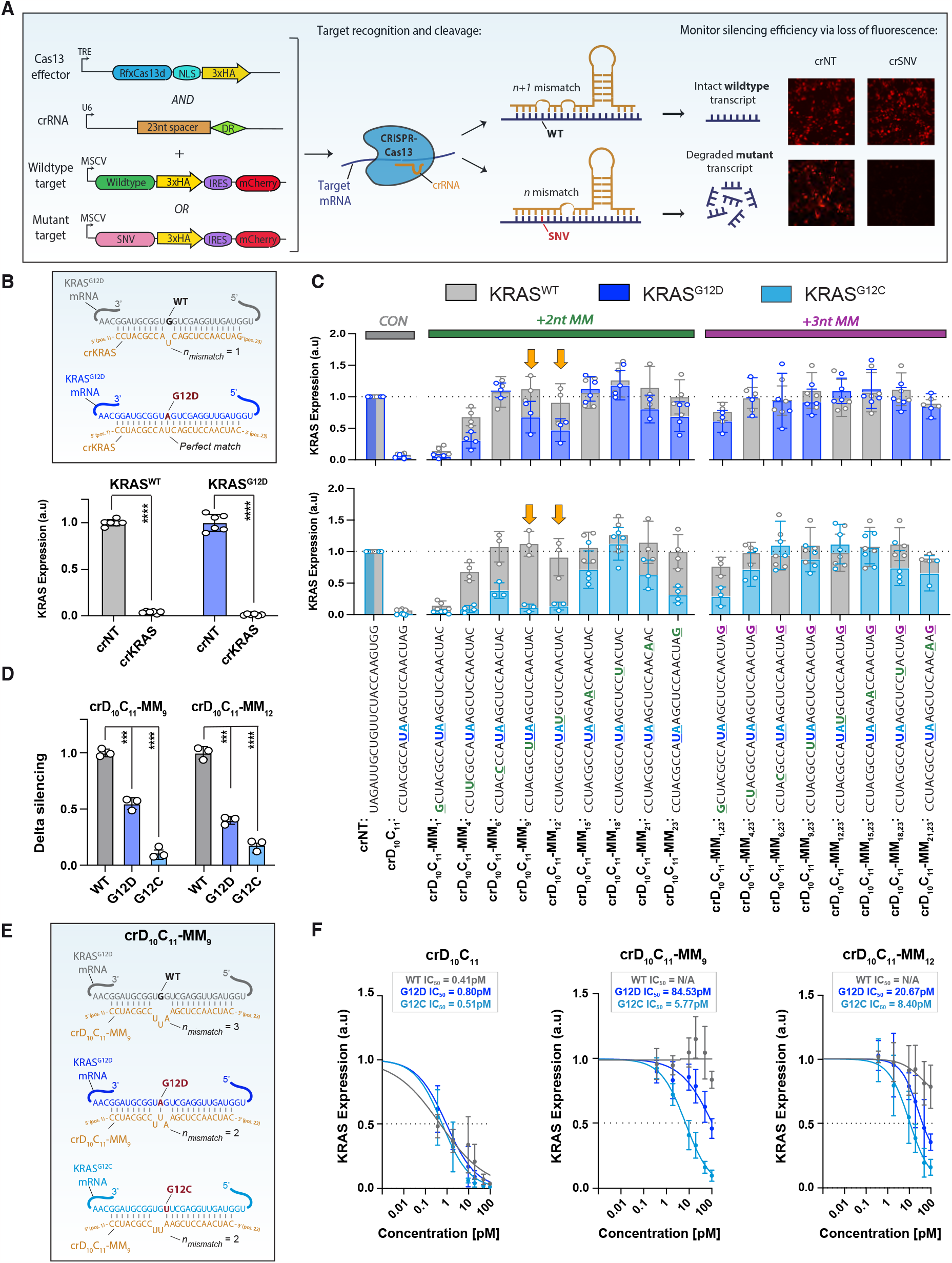
Selective cleavage of KRAS G12C and G12D mutant transcripts is achieved via precise crRNA mutagenesis. **(A)** Schematic of the RfxCas13d fluores-cence reporter assay used to assess silencing efficiency of wildtype and mutant tumour transcripts. **(B)** Schematic depicting the sequence and basepairing configuration of perfect-match crKRAS with WT KRAS versus G12D-mutant mRNA targets (upper). The silencing efficiency of crKRAS at 48h post-knock-in of WT (grey) or G12D-mutant (blue) KRAS, normalised against a non-targeting control crRNA (crNT; lower). **(C)** Systematic mutagenesis of non-selective crC10D11 was used to engineer G12D- and G12C-selective crRNAs. Sequences for each crRNA are shown below the barplot: the “U” nucleotide (in dark blue) indicates the uracil in the spacer sequence that basepairs with the G12D target and the “A” nucleotide (in light blue) indicates the adenine in the spacer sequence that basepairs with the G12C target (neither of which will basepair with WT target); the coloured nucleotides show the position of an additional mismatches introduced at various spacer locations through mutagenesis. Barplot shows silencing efficiency of crC_10_D_11_, and its mutagenesis derivatives against KRAS-WT (grey bars) vs KRAS G12D (dark blue bars; upper) or KRAS G12C (light blue bars; lower), normalised against a non-targeting control (crNT), at 48h post-transfection. crC10D11-MM9 and crC_10_D_11_-MM_12_ (orange arrows) show the highest selectivity against the G12 mutants. **(D)** Delta silencing efficiencies between WT (grey), G12D (dark blue) and G12C (light blue) KRAS variants for the top-performing crRNAs at 48h post-knock-in, indicating the degree of SNV-specificity. **(E)** Schematic depicting the sequence and basepairing configuration of SNV-selective crC_10_D_11_-MM_9_ with WT vs G12D- and G12C-mutant KRAS mRNA targets. **(F)** Dose-response curves derived from titration of parental crC_10_D_11_, crC_10_D_11_-MM_9_ and crC_10_D_11_-MM_12_ against KRAS WT (grey), G12D (dark blue) and G12C (light blue). All experiments were performed in HEK293T overexpression models. Individual data points show normalised fluorescence intensity averaged from four representative fields of view (from at least n=3 independent experiments), and error bars show mean ± SD. Statistical significance was determined using unpaired t-tests, where * p < 0.05, ** p < 0.01, *** p < 0.001, **** p < 0.0001.

Given that the position and number of nucleotide mismatches in crRNA spacer sequences can modulate silencing efficiency^23^,^29^ we sought to determine the optimal mismatch tolerance threshold that would confer preferential silencing of the G12-mutant transcript. As any additional perturbations to the crRNA sequence would be made in addition to the original SNV substitution, the number of mismatches in the wildtype sequence is always *n+l*, where *n* is the number of mismatches in the spacer sequence when using the SNV target as a template **(Figure lA)**. We first screened a series of RfxCas13d-compatible single-mismatch guides for selective silencing of KRAS G12D relative to KRAS WT.

Here, the position of the mismatch in each guide was shifted by one nucleotide, allowing the assessment of how each position of the 23-nt spacer sequence contributes to selective silencing activity. However, 22/23 mutagenesis derivatives achieved equipotent, efficient silencing of both variants **(Figure SlA)**, indicating that RfxCas13d can intrinsically tolerate single-nucleotide mismatches (MM). Whilst the crRNA with a mismatch at position 23 showed strongly selective silencing of KRAS G12D, indicating that this position may be important for achieving SNV-selectivity, the off-target silencing of KRAS WT was approximately 50% **(Figure SlA)**.

Thus, to maximise our ability to discriminate between WT and mutant KRAS transcripts, we performed systematic mutagenesis of this crRNA by introducing 2-3 additional synthetic mismatches along the spacer sequence, prioritising mismatches at nucleotide position 23 **(Figure lC)**. When choosing the position of the other synthetic mismatches in this crRNA series, we strategically noted that the KRAS G12D and G12C variants occur at adjacent nucleotides in the KRAS mRNA sequence (at positions c.34 and c.35, respectively; **Figure SlB)**. We leveraged this observation by ensuring that all mutagenesis derivatives retained the “U” nucleotide (corresponding to the G12D mutation) at crRNA position 10, but additionally incorporated an “A” nucleotide (matching G12C SNV) at crRNA position 11. Thus, by conserving this “D_1_0C_11_” motif across all guides, we anticipated that our engineered crRNAs would now be able to silence both G12D *and* G12C.

When screened against the G12D variant, all triple mismatch guides inefficiently silenced both WT and G12D **(Figure lC, upper)**, indicating that three mismatches in the crRNA may be the upper limit for efficient RfxCas13d mediated KRAS silencing. Conversely, certain double mismatch configurations conferred preferential silencing activity against the G12D variant **(Figure lC, upper)**. These SNV-selective guides contained synthetic mismatches at crRNA positions 9 (i.e., crD_1_0C_11_-MMg) and 12 (i.e., crD_1_0C_11_-MM_12_). Consistent with our G12D results, most triple-mismatch guides also showed poor on-target silencing against the G12C variant **(Figure lC, lower)**. Unlike G12D, crD10C1_1_-MM_2_3 exhibited SNV selective silencing of G12C, aligning with the results of our single-mismatch screen **(Figure lC, Figure SlA)**. Unexpectedly, the SNV-selectivity of crD_1_0C_11_-MMg and crD_1_0C_11_-MM_12_ was significantly enhanced when targeting G12C **(Figure lC, lower, Figure SlC)**, indicating that specific nucleotide that is mutated in each G12 variant (i.e., G>A for G12D vs. G>T for G12C) influences the silencing efficiency of individual crRNAs. When normalised against the WT variant, these two top performing guides showed significantly selective silencing of G12D (mean silencing 54.8% and 39.9%, respectively) and G12C (mean silencing 11.0% and 17.9%, respectively), respectively **(Figure 1D)**. This differential silencing activity is likely attributable to three mismatches with wildtype KRAS - exceeding the upper limit of mismatches that allow for efficient silencing - but only two mismatches with the G12D and G12C variants **(Figure lE, Figure S1D)**. Titration of these engineered crRNAs confirmed that this preferential silencing of G12C and G12D mutant transcripts was dose-dependent, with the half maximum inhibitory dose (ICso) estimated between 5-84pM and 8-21pM for crD10C11-MM9 and crD10C11- MM12, respectively. These SNV-selective crRNAs exhibited limited activity against WT KRAS at even the highest concentration of crRNA tested (ICsos non converged; **Figure lF)**. This highlights the tunability of this targeting strategy, as the optimal concentration of crRNA required to maximise silencing of the SNV transcript whilst also retaining minimal off-target silencing of the WT transcript can be easily identified via titration.

Collectively, these results indicated that directed mutagenesis of crRNA spacer sequences can unlock SNV selectivity.

### Exploiting mismatch tolerance enables selective targeting of all KRAS G12 hotspot mutations

Having confirmed the G12-selectivity of the crD10C11-MM9 and crD10C11-MM12 guides, we next questioned whether they would display cross-reactive silencing activity against the other c.34 and c.35 mutant KRAS variants **(Figure 2A)**. Beyond clinical relevance of Pan-RAS inhibition, targeting the full suite of G12 mutants also offers a unique opportunity to validate the relative importance of (i) the type of nucleotide substitution (i.e., G>A vs G>T) and (ii) the position of the SNV in the mRNA sequence (i.e., c.34 vs c.35) for efficient R/xCas13d mediated silencing. We therefore generated a library of KRAS G12 reporter plasmids encoding each of the remaining G12 hotspot mutations **(Figure S2A)** and assessed their sensitivity to crD10C11-MM9 and crD10C11-MM12. In contrast with the high on-target activity observed against G12C and G12D mutant transcripts, these crRNAs did not show efficient silencing of of the other c.34 (i.e., G12R, G12S) and c.35 (i.e., G12A, G12V) variants **(Figure 2B, S2B)**.

**Figure 2.**
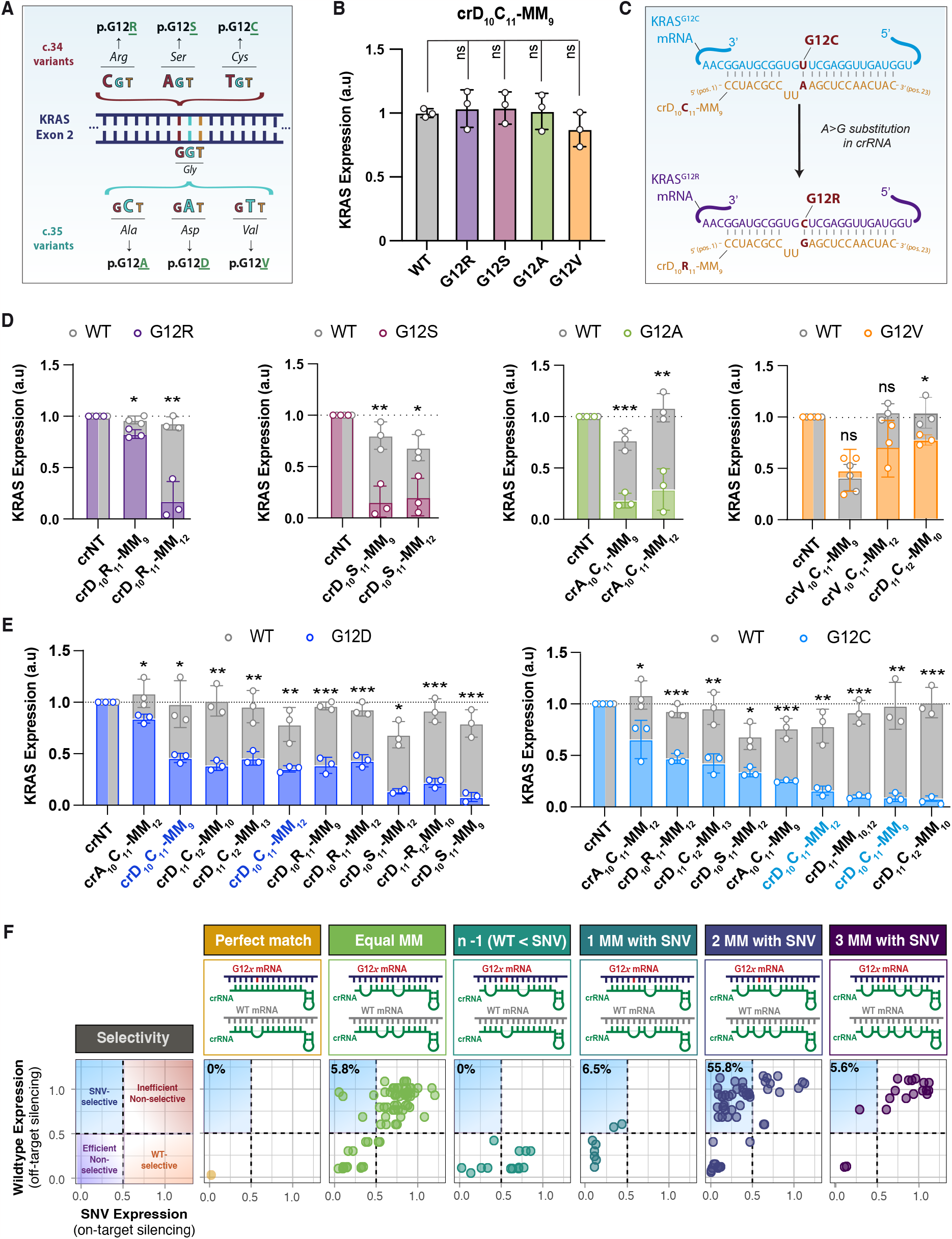
Exploiting the mismatch tolerance of RfxCas13d enables selective targeting of KRAS G12 hotspot mutations. **(A)** The G12 mutation hotspot is found in exon 2 of the KRAS gene (codon 12, nucleotides 34-36). For wildtype KRAS, the consensus coding sequence of “GGT” at codon 12 encodes a glycine (ie. G12). Missense mutations that affect the “G” nucleotide at position 34, collectively referred to as “c.34 variants”, change the amino acid sequence such that arginine (G12R, from c.34G>C substitution), serine (G12S, c.34G>A) or cysteine (G12C, c.34C>T) are encoded instead of glycine. “c.35 variants” arise from missense substitutions at nucleotide 35, causing glycine to be replaced by alanine (G12A, c.35G>C), aspartate (G12D, c.35G>A) or valine (G12V, c.35G>T). **(B)** Barplot showing minimal cross-reactive silencing by crC10D11-MM9 against other c.34 and c.35 KRAS variants (G12R, purple; G12S, pink; G12A, green; and G12V, gold), normalised against the silencing of WT KRAS, at 48h post-transfection. **(C)** Schematic depicting the mutagenesis strategy used to “switch” the silencing selectivity of our G12C- or G12D-selective crRNAs onto other G12 variants (G12R, G12S, G12A and G12V). In the example, the “A” nucleotide of the crC_10_D_11_-MM_9_ spacer, which basepairs with the “T” nucleotide present in G12C-mutated KRAS mRNA, can be switched to a “G” nucleotide, thus promoting basepairing with the “C” nucleotide present in G12R-mutated KRAS mRNA. **(D E)** Barplots showing silencing efficiency of various crC_10_D_11_-MM_9_ and crC_10_D_11_-MM_12_ mutagenesis derivatives against all KRAS G12 variants, normalised against crNT. Statistics indicate differential silencing efficiency between WT vs G12-mutants (rather than targeting vs crNT guide per variant). In (E), the parental crC_10_D_11_-MM_9_ and crC_10_D_11_-MM_12_ are labelled in bold. **(F)** Biplots showing the SNV-selectivity profile of all G12-targeting crRNAs in this study. Each point represents the mean silencing efficiency of a single guide against KRAS WT (y-axis) versus G12-mutant (x-axis) argets (from n=3 independent experiments). crRNAs that fall within the upper-left quadrant (light blue; indicating <50% expression of the G12 variant whilst maintaining >50% expression of the WT) are considered SNV-selective silencers. All experiments were performed in HEK293T overexpression models. For all bargraphs, error bars represent mean ± SD from three independent experiments. Statistical significance was determined using unpaired t-tests, where * p < 0.05, ** p < 0.01, *** p < 0.001, **** p < 0.0001.

We reasoned that specificity of our crRNAs could be “switched” from one variant to another by substituting the appropriate nucleotide at the c.34 or c.35 positions in the crRNA spacer **(Figure 2C)**. For example, a perfect match G12C-targeting guide may contain an “A” nucleotide in spacer position 11 (i.e., crC11), which will hybridise with the ‘‘T’’ nucleotide substitution found in G12C-mutated KRAS mRNA (i.e., c.34G>T). We reasoned that if we exchange the spacer nucleotide “A” for “G” at position 11 (i.e., crR11), we would promote basepairing with the “C” substitution present in G12R-mutated KRAS (i.e., c.34G>C), silencing activity might correspondingly “switch” from G12C to G12R **(Figure 2C)**. Using this strategy, we designed a series of crRNAs which conserved the structure of crD10C11-MM9 or crD10C11-MM12 but altered the nucleotide identity at the spacer positions involved in c.34 and c.35 base pairing (e.g., crD10R11-MM9, crA10C11-MM12, and so on). As predicted, directed mutagenesis of these crRNAs resulted in potent, SNV selective silencing of KRAS G12R, G12S and G12A relative to KRAS WT **(Figure 2D)**.

Unlike the other variants, G12V did not follow the expected silencing pattern, with crV10C11-MM9 and crV10C11-MM12 showing similarly inefficient silencing activity against both the WT and G12V variants **(Figure 2D)**. This equipotent silencing trend was confirmed in a further panel of putative G12V-selective guides **(Figure S2C)**, where all guides silenced both the wildtype and G12V variants with outstanding efficiency. Unexpectedly, one crRNA (crD11C12-MM10) demonstrated selective silencing of KRAS G12V relative to WT KRAS **(Figure 2D)**, despite having 3 mismatches with both G12V and WT and its intended targets being G12C and G12D. As the magnitude of G12V variant selective silencing was minor compared to the best candidate crRNAs for the other G12 variants, we did not proceed with further validation of the G12V target. However, this surprising result caused us to question whether any of our newly engineered guides may out-perform the parental crD10C11-MM9 and crD10C11-MM12 in selectively silencing G12C or G12D. Screening the entire panel of guides against each G12 mutant **(Figure S3)** uncovered several additional guides with improved G12-selectivity as compared with the parental guides **(Figure 2E)**, most notably for the G12D variant.

The broad variance in silencing efficiency of our crRNAs observed across a single G12 target (i.e., some G12C-targeting crRNAs are extremely efficient, and others less so) prompted us to investigate whether there are any generalisable features that could differentiate efficient SNV-selective versus inefficient or non-selective crRNAs. To that end, we pooled the silencing data from all crRNAs that targeted any KRAS variant and plotted their silencing efficiency relative to the number of mismatches (with a single G12 target) in their spacer sequences **(Figure 2F)**. As expected, the perfect-match crRNA for KRAS G12D (i.e., no mismatches with G12D, and one mismatch with KRAS-WT) efficiently silenced both WT and SNV with equipotency **(Figure 2F, yellow header)**. Similarly, when there is an equal number of mismatches in the spacer sequence for both the WT and the SNV transcript (e.g., 2 mismatches with WT but also 2 mismatches with the SNV variant), crRNAs generally exhibit no selectivity and typically silence both WT or SNV with equivalent efficiency (bottom-left quadrant) or inefficiency (top-right quadrant) **(Figure 2F, light green header)**. Predictably, when crRNAs are engineered using WT KRAS as template (instead of the SNV), thus meaning that the guide will have one *fewer* mismatch with the WT relative to the SNV (i.e., n-1, instead of n+l described previously), silencing efficiency is skewed in favour of the WT variant **(Figure 2F, dark green header)**.

Guides that contain one mismatch with the SNV (and two with the wildtype) are comparably efficient at silencing both WT and G12 variant **(Figure 2F, teal header)**, just as those with three mismatches with the SNV (and four with the wildtype) are comparably inefficient **(Figure 2F, purple header)**. Interestingly, over half of all crRNAs containing two mismatches with the SNV (and three with the wildtype) exhibit SNV-selective silencing **(Figure 2F, blue header)**, suggesting that this number of mismatches might be exquisitely important in the context of targeting mutant G12 sequences. However, the observation that certain guides with two nucleotide mismatches (relative to the SNV template) remain non-selectively efficient or inefficient suggests that the position of those two nucleotide mismatches may also influence silencing efficiency.

These data collectively suggest that intrinsic mismatch tolerance ‘rules’, related to the number and position of spacer mismatches, regulate the capacity of a crRNA to achieve $NV-selective silencing. Overall, guide RNAs that harbour two mismatches with the SNV and three mismatches with wildtype can achieve selective silencing of the SNV.

### Reprogrammed Cas13 represses ‘undruggable’ RAS mutants

Through our various KRAS targeting screens, we identified several crRNAs that exhibited strong preferential silencing of each G12 variant. For further validation, we selected the most potent selective silencers of G12A (crG12A), G12C (crG12C), G12R (crG12R), G12D (crG12D), and G12S (crG12S; **Figure 3A)**. For each guide, heteroduplex crRNA:mRNA formation was favoured in the mutant-targeting condition (lower t.G), consistent with the idea that increasing the number of spacer mismatches reduces binding affinity **(Figure 3A)**. Titration of these top-performing crRNAs confirmed the strikingly SNV-selective, dose-dependent silencing activity of these crRNAs against all G12 variants tested **(Figure 38)**. These guides were extremely potent, with ICs_o_s for each guide ranging from 1-22pM of plasmid. Conversely, the ICs_o_s for the WT could not be accurately calculated due to the failure of these guides to silence this variant with at least 50% efficiency. Moreover, G12-selectivity was maintained at the protein level for all variants, with silencing efficiency against the mutant protein enhanced by approximately 3- to 12-fold versus the wildtype protein **{Figure 3C-D)**. Collectively, these data show that reprogramming *RfxCas13d* via directed crRNA mutagenesis allows for strikingly selective inhibition of KRAS G12 hotspot mutants.

**Figure 3.**
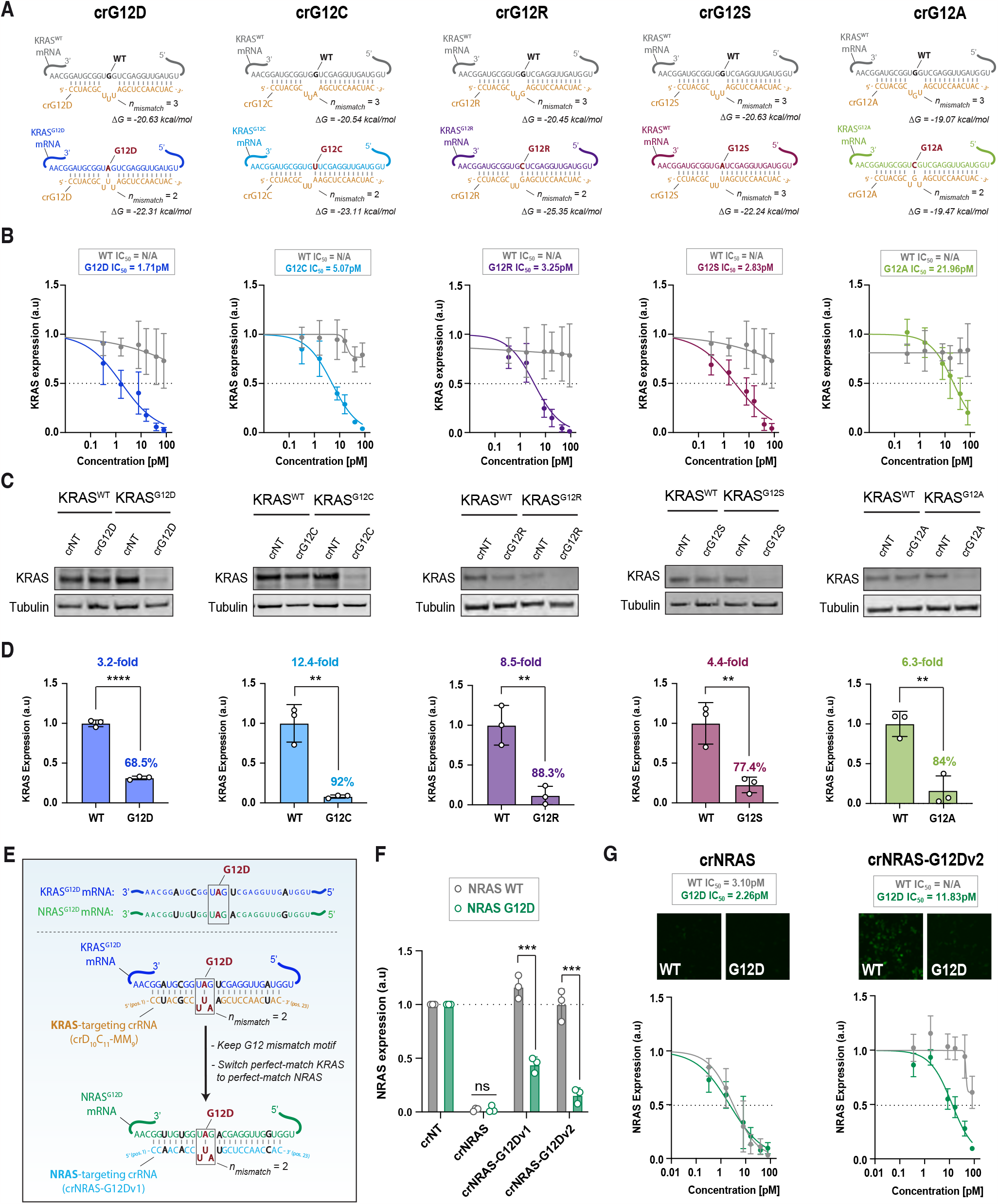
Potent, selective silencing of five KRAS G12 variants using reprogrammed RfxCas13d. **(A)** Schematic depicting the sequence and basepairing configuration of five mismatched crRNAs with wildtype KRAS versus G12-mutant mRNA targets. The number of mismatches (n) and change in Gibbs free energy upon heteroduplex formation (G) are shown beneath each crRNA:mRNA pair. **(B)** Dose-response curves derived from titration of top-performing crG12 guides against KRAS-WT vs a KRAS-G12 variant (-A, -C, -D, -R or -S). **(C)** Silencing efficiency of each guide assessed by WB in HEK293T cells transfected with KRAS WT or G12-mutated constructs. **(D)** Densitometry quantification of (C), showing SNV-specificity for all crRNAs at the protein level. **(E)** Schematic of the guide design strategy used to adapt KRAS-G12D targeting crRNAs to NRAS-G12D targeting crRNAs, showing sequence homology between the two mRNAs (upper) the sequence of each crRNA (lower). The box indicates the nucleotides surrounding the codon 12 hotspot, and nucleotides in black indicate bases which are not conserved between KRAS and NRAS wildtype sequences. **(F)** Silencing efficiency of parental crNRAS and SNV-selective crNRAS-G12Dv1 and crNRAS-G12Dv2 against NRAS WT (grey) and NRAS G12D (dark green), normalised against crNT. Individual data points show normalised fluorescence intensity averaged from four representative fields of view (from at least n=3 independent experiments). **(G)** Dose-response curves derived from titration of parental crNRAS and crNRAS-G12Dv2, against NRAS WT (grey) versus G12D (dark green). Error bars represent the mean fluorescence intensity ± SEM from eight fields-of-view. All experiments were performed in HEK293T overexpression models. Where statistics are shown, error bars indicate mean ± SD (or ± SEM in panel G) from at least n=3 independent experiments. Statistical significance was determined using unpaired t-tests, where * p < 0.05, ** p < 0.01, *** p < 0.001, **** p < 0.0001.

We next interrogated whether our strategy for targeting SNV could be extended to silence other prevalent oncogenic point mutations. To demonstrate this concept, we focussed on another member of the RAS family, NRAS, for which there are no targeted therapies. Single-base substitutions in NRAS are present in ∼10-15% of acute myeloid leukemias (AML), ∼40% of which are G12D SNVs (c.35 T>A) ^30^. The binding site of RfxCas13d surrounding the KRAS G12D motif differs by four nucleotides from the corresponding region in NRAS G12D **(Figure 3E, Figure S4A)**. Consequently, the specific crRNAs designed for KRAS G12D are not capable of silencing the NRAS G12D variant due to the presence of these four nucleotide differences, exceeding the mismatch tolerance threshold of RfxCas13d. We hypothesised that the KRAS G12D-specific crRNA could be modified into an NRAS G12D-specific crRNA by introducing targeted substitutions for these four differing nucleotides within the spacer sequence.

As expected, a crRNA that fully basepairs with the NRAS G12D binding site achieved extremely high silencing efficiency against both G12D and wildtype NRAS mRNA without selectivity. However, the two crRNAs we engineered to silence NRAS G12D (crNRAS-G12Dvl and crNRAS-G12Dv2, which adopt the G12-targeting motifs from KRAS-targeting crD10C11-MMg and crD10C11-MM12, respectively; **Figure 3E, Figure S4)**, showed clear G12D selectivity **(Figure 3F, S48)**. Consistent with our previous observations, this preferential NRAS G12D silencing effect was dose-dependent (G12D relative ICs_o_ = 11.83pM; **Figure 3G, S48)**. This data further underscores the effectiveness and adaptability of our strategy for targeting a broad range of oncogenic SNVs.

### SNV-selective silencing is achievable using orthogonal PspCas13b

To gain a deeper understanding of mismatch tolerance and SNV-specific silencing rules, we expanded our investigation to additional point-mutated transcripts and Cas13 orthologues. Specifically, we questioned whether SNV-selective silencing is unique to *RfxCas13d* or whether this property is instead a conserved feature across Cas13 family members. . For this, we investigated whether another Cas13 orthologue, *PspCas13b*, could also achieve SNV-selective silencing. Of note, *PspCas13b* possesses a 3O-nt long spacer sequence (versus 23-nt for RfxCas13d) which may confer enhanced target specificity but also a higher mismatch tolerance.

To show that single-base resolution targeting can silence other onco-transcripts (which have limited sequence homology with KRAS), we focussed our PspCas13b designs on targeting the BRAF^vGooE^ mutation, in which a single nucleotide substitution (c.1799T>A) results in the replacement of valine by glutamate at amino acid position 600. This SNV is the most common BRAF aberration and is found in approximately 7% of all human cancers and up to 60% of melanomas 31. We first designed four *PspCas13b* perfect-match crRNAs (crBRAFl-4) which tile across the V6OOE point mutation.

These guides incorporated “GG” in spacer positions 1-2, in line with our recent report that this motif leads to enhanced PspCas13b-mediated silencing ^23^• Using BRAF reporter assays, all perfect-match guides enabled potent BRAF mRNA cleavage relative to the non-targeting control, among which the crBRAF-1 guide, which basepairs with V600E at spacer position 5, showed the highest silencing activity **(Figure 4A, Figure SSA)**.

**Figure 4.**
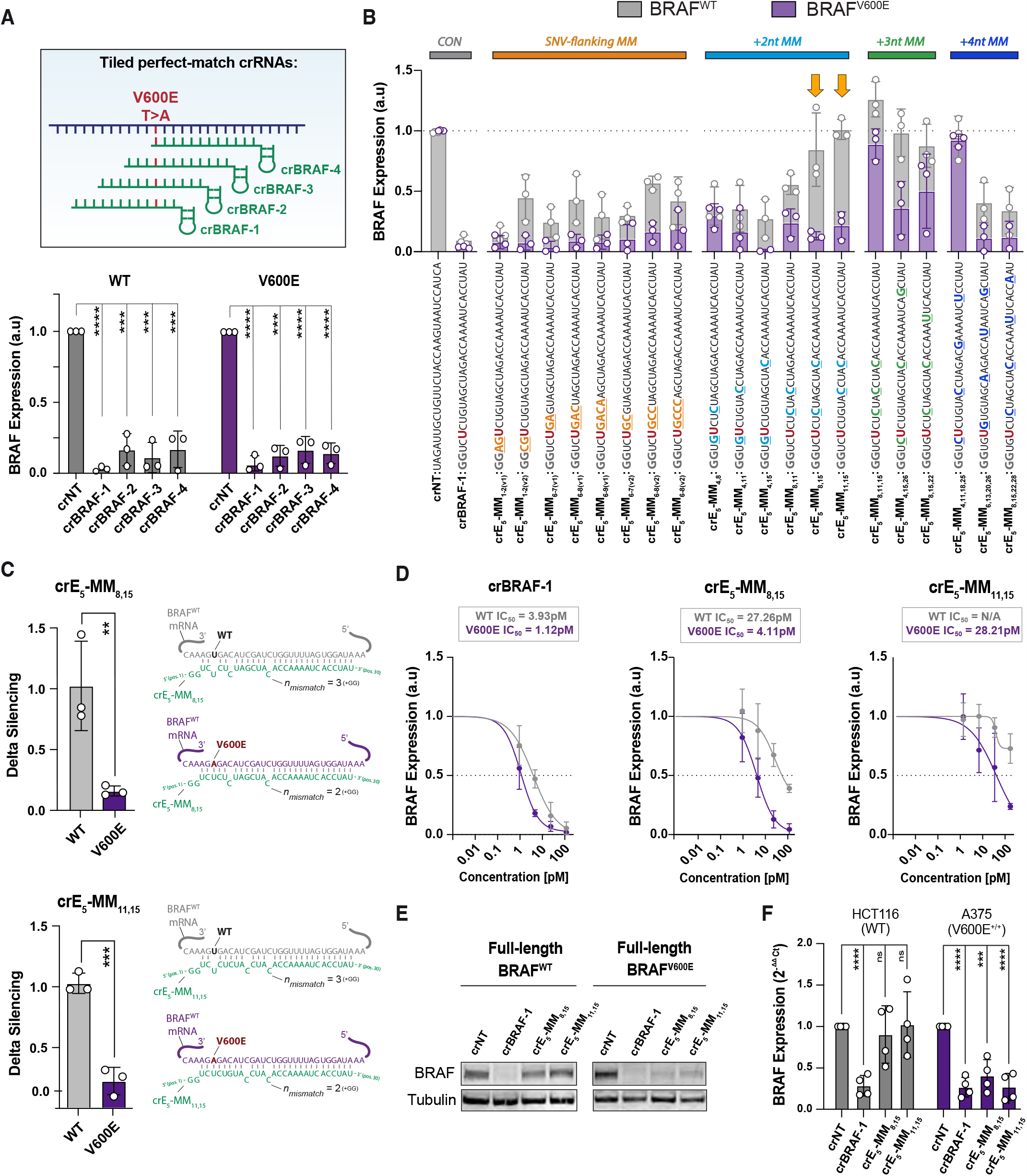
SNV selective silencing of BRAF V600E using orthogonal PspCas13b. **(A)** Silencing efficiency of four perfect-match, BRAF-targeting crRNAs at 48h post-knock-in of WT (grey) or V600E (purple BRAF variants, normalised against crNT. **(B)** Systematic mutagenesis of non-selective crBRAF-1 was used to engineer V600E-selective crRNAs. Sequences for each crRNA are shown below the barplot; the “U” nucleotide (in red) indicates the uracil in the spacer sequence that basepairs with the target V600E but not with the WT BRAF RNA, and the coloured nucleotides show the position of an additional mismatch introduced at various spacer locations through mutagenesis. Barplot shows silencing efficiency of crBRAF-1 and its mutagenesis derivatives against BRAF-WT (grey) vs BRAF-V600E (purple), normalised against crNT at 48h post-transfection. crE_5_-MM_8,15_ and crE_5_-MM_11,15_ (orange arrows) show the highest selectivity against BRAF V600E. **(C)** Delta silencing efficiencies between WT (grey) and V600E (purple) variants for the top-performing crRNAs, indicating the degree of SNV-specificity. Schematics depicting the sequence and basepairing configuration of crE5-MM_8,15_ and crE5-MM_11,15_ with WT vs V600E-mutant BRAF mRNA targets are shown adjacent. **(D)** Dose-response curves derived from titration of parental crBRAF-1, crE_5_-MM_8,15_ and crE5-MM_11,15_ against WT (grey) or V600E (purple) BRAF at 48h post-transfection. For all graphs in A-C, individual data points show averaged fluorescence intensity from eight representative fields of view (n=3 independent experiments), and error bars show mean ± SD. **(E)** Silencing efficiency assessed by WB in HEK293T cells transfected with PspCas13b and full-length WT or SNV constructs. **(F)** Silencing efficiency assessed by RT-qPCR in cancer cell lines (A375 melanoma and HCT116 colorectal cancer) which endogenously express the SNV-containing oncogene (mean ± SD from n=3 independent experiments). Statistical significance was determined using unpaired t-tests, where * p < 0.05, ** p < 0.01, *** p < 0.001, **** p < 0.0001.

However, none of these guides were able to discriminate between wildtype and SNV-containing targets.

To achieve SNV-selectivity, we serially mutated crBRAF-1 by introducing either contiguous blocks of 2-4 mismatched nucleotides flanking the SNV site, or a series of two, three, or four single-nucleotide mismatches that were distributed along the spacer sequence **(Figure 4B)**. Whilst many crRNAs retained their ability to silence the mutant transcript, often with efficiencies comparable with the parental perfect-match crBRAF-1 (e.g. crE_5_-MM_14,15_)., certain crRNAs completely lost their silencing activity (e.g. crE_5_-MM_4,11,18,25_). As expected, silencing activity against the BRAF wildtype target was more severely impaired as these transcripts had one additional mismatch (n+l) for each of the spacer sequence screened. The crRNAs that were least efficient at silencing wildtype BRAF transcripts were over-represented in the mutagenesis groups where spacers had two- or three mismatches in the VG00E spacer (corresponding to three and four mismatches with the wildtype, respectively), suggesting that this may represent an optimal mismatch tolerance threshold for the PspCas13b orthologue when targeting BRAF VG00E sequence. Of these, two crRNAs, crE_5_-MM,_8,15_ and crE_5_-MM_11,15_, showed significant differential **(Figure 4C, SSB)** and dose-dependent **(Figure 4D)** silencing activity in favour of the VG00E transcript.

As the BRAF construct utilised in our reporter assay encoded a truncated, VG00E-spanning BRAF gene (∼soont), we next sought to investigate whether the SNV specificity of crVG00E-13 and crVG00E-14 would be retained when targeting the full-length BRAF transcript. HEK293T cells transfected with constructs encoding full length BRAF-WT or BRAF^v6ooE^ also exhibited the expected pattern of silencing, with both guides preferentially knocking down BRAF^v6ooE^ at the protein level **(Figure 4E)**. This indicates that the silencing efficiency of these pre validated crRNAs was not disrupted by any potential secondary structures present in the full-length transcripts expressed in HEK 293T cells.

Next, we tested the silencing activity of the engineered crRNAs in cancer cell lines with different BRAF genotypes, HCT116 (colorectal cancer, WT) and A375 (melanoma, homozygous VG00E). Corroborating the results seen in HEK293T cells **(Figure SSB)**, this data confirmed that the efficiency and SNV-selectivity of these crRNAs are retained in cancer cell lines expressing endogenous BRAF variants **(Figure 4F)**. Collectively, these data indicate that the *PspCas13b* system can be reprogrammed to target VG00E-mutated transcripts in both overexpression and endogenous contexts with high specificity. Moreover, our demonstration of SNV selective silencing in two orthogonal Cas13 systems reinforces both the sensitivity and versatility of our screening platform to identify SNV-selective crRNAs and suggests that it may be applicable to other SNVs beyond **KRAS** G12 or BRAF VG00E.

## DISCUSSION

In this study, we developed single-nucleotide precision Cas13 tools capable of selectively degrading various oncogenic SNV transcripts with high efficiency and very limited off-targeting of wildtype mRNA. The engineering required for this single-base precision RNA targeting was challenging due to the intrinsic mismatch tolerance of various CRISPR-Cas13 enzymes^18,23^. Directed perturbation of spacer-target interactions at precise spacer nucleotide positions through mutagenesis enabled us to identify key positions that are highly sensitive to mismatches, facilitating *de novo* crRNA design that can discriminate between point-mutated transcripts and their wildtype variants. Beyond the oncogenic KRAS, NRAS, and BRAF SNV transcripts we silenced in this study, we aimed to develop a systematic approach to predicting personalised single-base precision crRNAs capable of targeting other pathogenic SNVs. This strategy was previously unattainable due to the limited comprehension of the molecular basis of mismatch tolerance and intolerance at single nucleotide resolution.

Consistent with previous reports^32^, different regions of the *RfxCas13d* spacer showed significant variation in mismatch intolerance, with the highest mismatch sensitivity exhibited in the central region. When we interrogated generalisable *RfxCas13d* design principles to enable selective silencing of KRAS SNVs, we found that the transition from mismatch tolerance (active nuclease) to intolerance (inactive nuclease) often occurs when three nucleotide mismatches are introduced at the central region of *RfxCas13d’s* spacer-target duplex. Two nucleotide mismatches with point-mutated transcripts were well-tolerated and resulted in potent silencing of SNVs, whereas the additional third mismatch with the wildtype transcript led to a significant or complete loss of silencing. We speculate that this mismatch intolerance threshold (three nucleotides) may vary when targeting other sequences due to the thermodynamic stability of the A-form RNA helix in the enzyme’s central cleft and the potential for subtle, target-specific interactions between the duplex RNA and the surrounding amino-acid residues of *RfxCas13d* that are intrinsic to the target sequence. Indeed, other high throughput studies using different spacer-target sequences have shown that the sensitivity to a single, double, or triple nucleotide mismatches may be dependent on the target sequence^25^.

Our results indicated that a single additional mismatch often confers a sudden transition of Cas13 from an active to inactive state. Although the predicted change in Gibbs free energy favours the hybridisation of a given crRNA with the mutant mRNA sequence (i.e. lower tiG for crRNA:mutant mRNAs heterodimers versus crRNA:wildtype mRNAs; **Figure 3)**, it is unlikely that this modest reduction in binding affinity alone can explain the profound shift from mismatch tolerance to intolerance. Rather, the mismatch intolerance threshold may be crossed due to steric hindrance created by bulky unpaired nucleotides at the central cleft of the enzyme, which may constrain the molecular rearrangement required for HEPNl and HEPN2 nuclease domain activation^**33**,^ Indeed, structural studies of the Cas13d-crRNA binary complex demonstrated that various regions of the spacer positioned within the open channel of the enzyme are exposed to the solvent^**33**,^ suggesting that the Cas13d spacer is unlikely to possess a unique mismatch-sensitive seed region responsible for initiating basepairing with the target. Instead, the sensitivity to mismatches in the central region we observed could be due to steric hindrance and the impediment of domain rearrangement. Future structural and dynamic studies may further illuminate the molecular components of this transition from mismatch tolerance to intolerance when various mismatches are located at the central region of the R/xCas13d spacer.

Interestingly, when we interrogated the single-base precision capability of the *PspCas13b* ortholog using a different target sequence (BRAF-V600E), we also found that mismatch number and position are critical for the activity of this Cas13 ortholog, although their number and position vary from those of R/xCas13d. crRNAs harbouring two non-consecutive mismatches with the mutant template (at position 15, combined with position 8 or 11, with an additional mismatch for the wildtype template arising from the SNV located at position 5) exhibited the most noticeable discrimination between wildtype and BRAF-V600E transcripts. This precise balance between the number and position of mismatches suggested the existence of asymmetry within the spacer with some regions highly sensitive to mismatches while others are resilient. We found that this mutagenesis strategy generated at least one crRNA capable of selective and potent silencing of each of the six possible KRAS G12 mutants **(Figure 3)**, albeit with a lower silencing potency for G12V compared to the other variants. Further development of these Cas13-based KRAS suppressors as RNA therapeutics could be of high clinical significance, as 90% of KRAS-driven cancers remain clinically unactionable ^**7**^**•**

In this proof-of-concept study, we have demonstrated a highly precise RNA targeting strategy which effectively suppresses single-nucleotide variants whilst sparing wildtype transcripts. This tool may be of great value for elucidating the function of point-mutated variants of unknown significance or, with further development, to therapeutically suppress known oncogenic drivers with single-nucleotide precision. Despite the significant potential, the major hurdle for these RNA-targeting tools lies in the absence of efficient and safe strategies for delivering large CRISPR enzymes and their crRNAs *in vivo*. Nonetheless, important milestones have been made in the delivery of various CRISPR systems using AAVs ^**34**,**35**^, or through the delivery of Cas endonuclease mRNA encapsulated into lipid nanoparticles ^**36-39**^These novel delivery modalities lay a strong foundation for future studies in patient-derived cell lines and animal models which are needed for the further advancement of these SNV-specific Cas13 drugs.

## ACKNOWLEDGMENTS

The authors thank all lab members from the Trapani, Ekert, and Fareh labs for facilitating experiments and discussions. We also thank A/Prof Ilia Voskoboinik and his lab members for other valuable discussions.

## AUTHOR CONTRIBUTIONS

M.F and C.S conceived the study. M.F and P.G.E supervised the study. C.S and M.F designed the experiments. C.S and R.Y performed the experiments and analysed the data. All authors discussed the project and the data. C.S and R.Y generated all graphs and figures. C.S and M.F wrote the manuscript. All authors read, commented, edited, and approved the manuscript.

## DATA AVAILABILITY

All data are available in the main text and/or supplementary materials. Source Data are provided with this paper. All key plasmids constructed in this study, their sequences, and maps will be deposited to Addgene upon publication.

## SUPPLEMENTARY DATA

Supplementary data are available online.

## MATERIALS AND METHODS

### Generation of SNV-selective crRNAs

#### crRNA design and cloning

RfxCas13d crRNA spacers were designed as 27-nt single stranded forward and reverse oligos containing AAAC and AAAA overhangs, respectively, allowing for ligation into BsmBI digested plasmid that encodes RfxCas13d direct repeat (Addgene #138150). PspCas13b crRNA spacers were designed as 34-nt single-stranded forward and reverse oligos containing CACC and CAAC overhangs, respectively, allowing for ligation into Bbsl-digested plasmid that encodes PspCas13b direct repeat (Addgene #103854). Oligos were ordered as single stranded DNA (IDT) at a concentration of 100 μM. All crRNA spacer sequences used in this study are available in Table Sl. To produce double-stranded DNA oligos suitable for plasmid ligation, 3 μM of each forward and reverse DNA oligos were annealed in T4 DNA Ligase buffer (NEB) by heating to 95° C for 5 mins then cooling at a rate of 4°C every 2 minutes until ambient temperature was reached. The PspCas13b crRNA backbone plasmid (Addgene #103854), a gift from Feng Zhang 40, was digested using Bbsl (NEB) in lx NEBuffer r2.1 at 37°C for 2 hours. The RfxCas13d crRNA backbone plasmid (Addgene #138150), a gift from Neville Sanjana ^26^, was digested using BsmBl-v2 (NEB) in lx NEBuffer r3.1 at 55°C for 2 hours. Digested plasmids were separated on a 1% (w/v) agarose gel in lx TAE buffer at 100V for 1hr. The linearised fragment was then excised and purified using the NucleoSpin Gel and PCR Clean up Kit (Macherey-Nagel). 400nM of the annealed oligos were ligated with lng of digested crRNA backbone using T4 ligase (Promega) in lx T4 ligase buffer for 3hrs at RT.

#### Nomenclature of SNV-se/ective crRNA

To ensure clarity, we have devised a consistent nomenclature system for crRNAs that effectively conveys information about the target single nucleotide variant (SNV) and the position of any introduced mismatches. Uppercase characters within the nomenclature signify the specific SNV being targeted by the respective crRNA. Additionally, the designation **‘MM’** denotes the presence of synthetic mismatches that have been introduced through spacer mutagenesis. The subscript numbers within the nomenclature correspond to the positions within the crRNA spacer, with position 1 situated at the 5’ end. For instance, considering the crRNA ‘crD10C11-MMg,^1^ which is designed to basepair with the KRAS G12D transcript. Here, the SNV interacts with spacer nucleotide 10. The same crRNA can also basepair with the KRAS G12C transcript, wherein the SNV interacts with spacer nucleotide 11. This crRNA forms an additional synthetic mismatch at spacer position 9, relevant to both KRAS G12D and KRAS G12C targets. A visual representation of this concept can be found in the schematic in Figure lE.

### Prediction of crRNA:mRNA heterodimer affinities

*1:,G* analyses were performed using the RNAcofold module of the Vienna RNA webserver 41 available at: http://rna.tbi.univie.ac.at/cgi-bin/RNAWebSuite/RNAcofold.cgi.

### Cloning of oncogene expression constructs

The BRAF and KRAS coding sequences were obtained from NCBI and all fragments for cloning were ordered as dsDNA gBlocks (IDT). Truncated BRAF inserts for cloning were designed to contain 250bp up- and downstream of codon V600, adding in frame reporter tags to the 3’ end (3xFLAG for BRAF-WT and 3xHA for BRAF-V600E), and flanking the excerpt with EcoRI/Xhol restriction cut sites. Full-length KRAS-WT was engineered in-frame with 3xHA and BamHI/Xhol cut sites. Cloning into MSCV-IRES reporter backbones encoding GFP (for BRAF-WT) or mCherry (for BRAF V600E and KRAS) was achieved via plasmid digestion, gel purification, and T4 ligation as previously described ^18^• Plasmids encoding full-length BRAF wildtype (p3xFLAG-CMV-BRAF, Addgene #131710) and V600E mutant (p3xFLAG-CMV-BRAF-V600E, Addgene #131723) were gifts from Jacques De Greve. (29)

### Generation of point mutated plasmids via site-directed mutagenesis

SNV-containing plasmid constructs for KRAS G12 (-A, -C, -D, -R, -S, -V) mutants were generated using site-directed mutagenesis. S0ng of parental plasmid was amplified with complementary 35nt-long forward and reverse primer pairs (0.25uM each) containing the desired nucleotide substitution (at nt position 17) in the presence of 1.25U *Pfu* DNA polymerase (Promega) and 200uM dNTPs (each) in lx MgSO4 buffer. After initial denaturation at 95°C, the reaction mix underwent 15 cycles of denaturation (95°C, 30s), annealing (58-60°C, 30 seconds) and extension (72°C, 2min/kB) followed by final extension (72°C, Smin). Parental plasmids were selectively linearised from the PCR product via digestion with Dpnl (Promega) in lx MULTI-CORE buffer for 1hr at 37°C and the digestion product was immediately transformed into Stbl3 competent bacteria.

### Plasmid amplification and purification

SμL of ligation product or lO0ng of circularised plasmid were transformed into 25μL of Stbl3 competent bacteria by heat shock at 42°C for 90 seconds, followed by 2min on ice. The transformed bacteria were recovered in 250μL of LB broth media for 1 hour at 37°C in a shaking incubator (200 rpm). The bacteria were pelleted by centrifugation at 8,000 rpm for 1 minute, resuspended in 30μL of LB broth and plated onto pre warmed LB agar plates containing 75μg/mL ampicillin. Following overnight incubation at 37°C, single colonies were picked and transferred into 3mL bacterial starter cultures and incubated for ∼16 hours prior to mini- or maxi-prep DNA purification using the NucleoSpin Plasmid kit (Machery Nagel). Successful cloning of all plasmids was verified by Sanger sequencing (Australian Genome Research Facility).

### Cell culture

HEK293T (ATCC CRL-3216) cells were cultured in DMEM high glucose media (Gibco). A375 (ATCC CRL-1619) and HCT-116 (ATCC CCL-247) were cultured in RPMI. Both media were supplemented with 10% heat inactivated foetal bovine serum (Life Technologies), 2mM Glutamax (Gibco), and 1% Penicillin/Streptomycin (Gibco). HEK293T cells were incubated at 37°C and 10% CO2. All other cell lines were incubated at 37°C and 5% CO2. Cells were passaged twice a week and routinely tested for mycoplasma.

### RNA silencing assays via transient transfection

This approach requires cloning the coding sequence of the KRAS single-nucleotide variant (SNV) of interest into an IRES fluorescent protein (FP) reporter construct (as described above). When transfected, this construct generates a chimeric SNV-IRES-FP transcript that is subsequently translated into two separate proteins due to the presence of the IRES sequence. Cas13 cleaves the SNV-IRES-FP chimeric RNA at the SNV site, resulting in RNA fragments lacking key features such as 5’ cap, 5’ and 3’ UTRs, and polyA tail, which leads to reduced intracellular RNA stability and/or compromised ribosomal translation. Consequently, the cleaved SNV fragment and downstream reporter gene undergo degradation and/or translational repression, resulting in loss of fluorescence signal. Therefore, efficient cleavage of the SNV transcript by Cas13 is accompanied by the loss of fluorescence signal.

Transfection experiments to determine the silencing efficiency of each crRNA were carried out using an optimised version of the manufacturer’s Lipofectamine-3000 protocol (lnvitrogen). Approximately 18h prior to transfection, HEK293T were seeded at 25,000 cells/100μL or 150,000 cells/S00μL densities in 96-well or 24-well densities in tissue culture treated flat-bottom plates (Corning), respectively. For each transfection condition, performed in duplicate, two master mixes were prepared. For 96-well plates, master mix A contained 25ng Cas13 effector plasmid (PspCas13b or RfxCas13d), target plasmid (S0ng BRAF- or 100ng KRAS expression plasmids), crRNA plasmid (0.1-100ng), and 4% (v/v) P3000 enhancer reagent {lnvitrogen) in Opti-MEM Serum-free Medium (Gibco), making up SμL total volume for 96-well plates. Master mix B contained 6% (v/v) Lipofectamine-3000 reagent {lnvitrogen) in Opti-MEM Serum-free Medium, making up SμL total volume. Master mixes A and B were combined at a 1:1 ratio and incubated at room temperature for 20 minutes, after which 10μL of A+B master mix was added to each well. For larger plates {24w, 6w), seeding density and all volumes were upscaled by surface area. For western blotting experiments shown in Figure 3C, the crRNA concentration was increased to 300ng per 24w. As the RfxCas13d effector plasmid is inducible, doxycycline at a final concentration of lug/ml was added to each well at the time of transfection. Transfected cells were incubated at 37°C in 10% CO_2_ for 48h, then imaged using the EVOS MS000 FL Cell Imaging System {ThermoScientific). The mean fluorescence intensity of each image was quantified using an *in-house* macro written for lmageJ software in batch mode. Briefly, images were converted to 8-bit, threshold adjusted based on the fluorescence intensity of positive (crNT) and negative (untransfected) controls, converted to a binary mask, and the fluorescence intensity per pixel measured using the Analyze Particles function. Mean fluorescence intensities were obtained from four fields of view per well for each crRNA and subsequently normalized to the crNT control. Plasmid relative ICS0s, where shown, were obtained by first calculating the molecular weight (MW) of the plasmid (DNA length in bp x 650, which is the average MW of a single DNA bp), then calculating the molarity (i.e. mass transfected per 110μL total volume per well of a 96w plate).

[Note: we found that that transfection of equivalent mass of the two full-length BRAF plasmids led to marked difference in BRAF protein expression, where V600E expression was several-fold higher than for BRAF-WT. Thus, for transfection experiments utilising these plasmids, we titrated the amount of each plasmid such that they produced equivalent protein expression measured by western blot. We balanced the overall total amount of DNA transfected in each condition by co-transfecting the residual amount of empty vector plasmid (of equivalent size).]

### Cell sorting, RNA extraction, cDNA synthesis, and RT-qPCR

A375 and HCT116 cells were sorted 48h post-transfection {for Cas13b-BFP+ cells) using a BD FACSAria Fusion 5 or Fusion 3 instrument. For all flow cytometry experiments, single-cell suspensions were resuspended in FACS Buffer {2% FBS, 0.2% EDTA in PBS) and passed through a 35 μm nylon mesh cell strainer to exclude large aggregates. Single cells were gated based on morphology and the threshold for BFP positivity was selected using an untransfected cell suspension as a non fluorescent control. Total RNA was isolated from the recovered cells (∼Sx10^5^ cells) using NucleoZOL one-phase purification (Machery Nagel) according to the manufacturer’s instructions. lμg of total RNA was converted to cDNA using the high capacity cDNA reverse transcription kit *(ThermoScientific)* following the manufacturer’s instructions. Quantitative RT-PCR reaction was performed in duplicate in a StepOne Real-Time PCR system *(ThermoScientific)* using PowerUp^™^ SVBR^™^ Green Master Mix *(ThermoScientific)*. The total reaction mixture contains 0.2μ1 cDNA, 0.6μM forward primer, and 0.6 **μM** reverse primer (sequences available in Table 52).

### Western blotting

Cells were washed with ice-cold PBS and lysed on ice in RIPA lysis buffer {SO mM Tris, 150 mM NaCl, 1% NP-40, 2% SDS, pH 7.5) containing protease inhibitor cocktail (Roche, 04693159001) and phosphatase inhibitor cocktail (Roche, 4906845001). Samples were incubated for 30min at 4 °C with rotation {25 rpm) and centrifuged at 16,000 g for 10 min at 4 °C, after which the supernatants were recovered. Protein concentrations were determined using the Pierce BCA protein assay kit (ThermoScientific) according to the manufacturer’s instructions. Samples for polyacrylamide gel electrophoresis were prepared by combining 25μg protein lysate with 4x Bolt LDS Sample Buffer {lnvitrogen) and 10x Bolt Sample Reducing Agent (lnvitrogen) in 30 μL total volume and denatured at 95°C for 5 minutes. Samples were loaded adjacent to a PageRuler prestained protein ladder (ThermoScientific) onto Bolt Bis-Tris 4-12% gels Mini Protein Gels (lnvitrogen) in lx MES SDS running buffer {lnvitrogen). Polyacrylamide gels were electrophoresed at 100 V for 1.5 hours. Following electrophoresis, proteins were transferred onto PVDF membranes (activated in 100% methanol for 5 seconds, rinsed with MilliQ H2O, then soaked in lx transfer buffer (Tris-HCI 25mM [pH7.3], glycine 192mM, 20% v/v methanol)) using a Trans-Blot SD Semi-Dry Transfer Cell (Bio-Rad, CA, USA) apparatus {20V, 30min). After protein transfer, membranes were blocked with 5% skim milk (w/v in PBS/0.5% tween [PBS T]) at RT for 1 hour, then incubated with primary antibody (Table 53) diluted in 5% (w/v) skim milk and incubated on a roller at 4°C overnight. The membrane was washed with PBS-T (3 x Sm) and incubated in secondary antibody (Table 53) diluted in Intercept Blocking Buffer (LI-COR) for 1 hour at RT. After washing (3 x Smin), the membrane was imaged using an Odyssey Dlx lmager (LI-COR). Densitometry quantification was performed using Image Studio software (LI-COR).

### Data and Statistical Analyses

All experiments were performed in at least 3 biological replicates unless otherwise indicated, and data is expressed as mean ± standard deviation (SD). For data presented in Figure 2F, silencing efficiencies against wildtype vs SNV were averaged from three independent experiments and plotted in R (version 4.1.0) using the ggplot2 package (script available in source data file). All statistical analyses were performed with GraphPad Prism 9 Software {GraphPad, CA, USA, v9.4.1), and t-tests were used to compare pairs of data. p<0.05 was used as the threshold of rejecting the null hypothesis, and statistical significance is shown as:*= p<0.05,**= p<0.01,***= p<0.001, ****= p<0.0001.

