## supplemental file for "Molecular principles of CRISPR-Cas13 mismatch intolerance enable selective silencing of point-mutated oncogenic RNA with single-base precision"

**Figure S1**

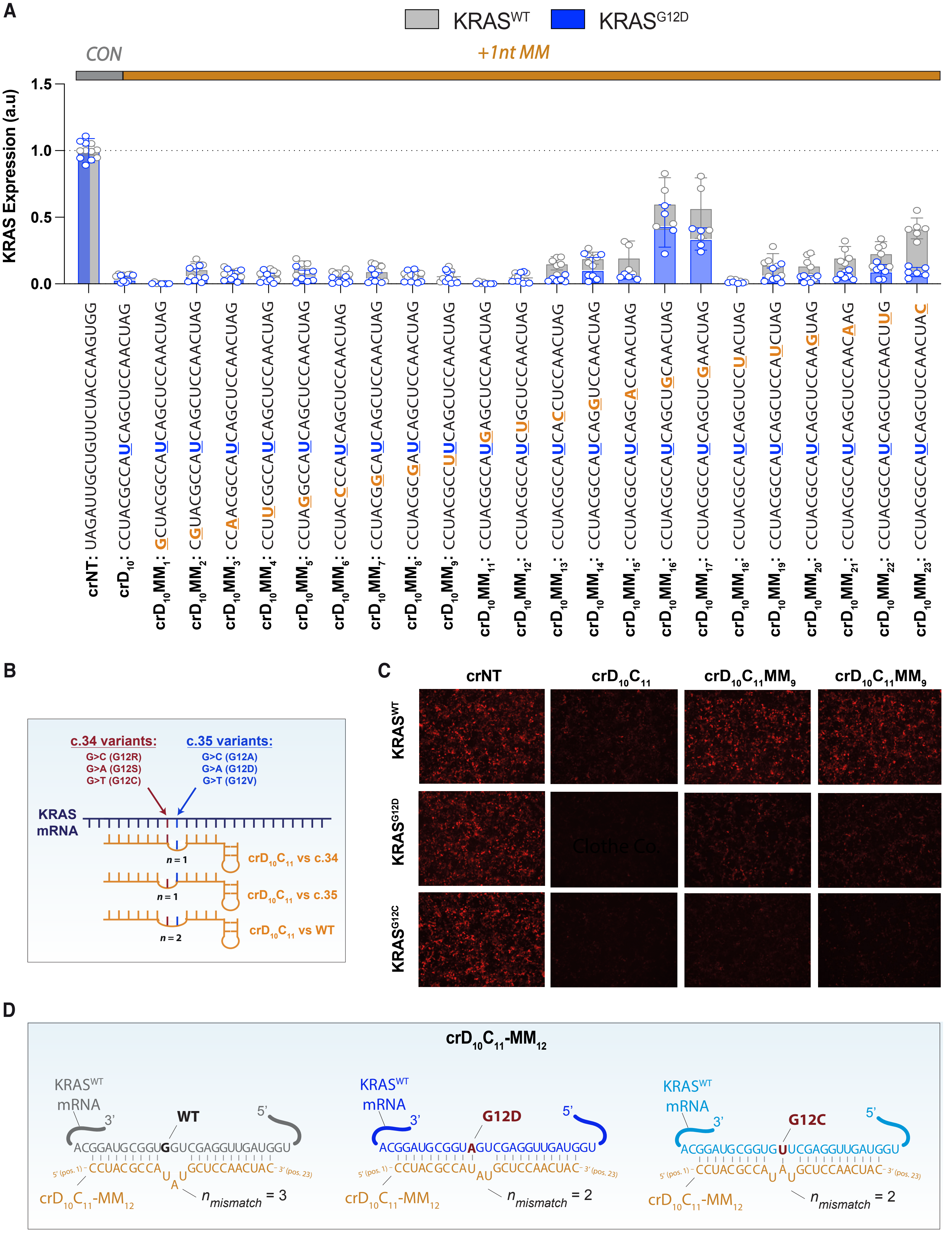

A
KRAS^WT
KRAS^G12D
CON
+1nt MM
KRAS Expression (a.u)
1.5
1.0
0.5
0.0
crNT: UAGAUUGCUGUUCUACCAAGUGG
crD10: CCUACGCCAUCAGCUCCAACUAG
crD10MM1: GCUACGCCAUCAGCUCCAACUAG
crD10MM2: CGUACGCCAUCAGCUCCAACUAG
crD10MM3: CCAACGCCAUCAGCUCCAACUAG
crD10MM4: CCUUCGCCAUCAGCUCCAACUAG
crD10MM5: CCUAGGCCAUCAGCUCCAACUAG
crD10MM6: CCUACCCCAUCAGCUCCAACUAG
crD10MM7: CCUACGGCAUCAGCUCCAACUAG
crD10MM8: CCUACGCGAUCAGCUCCAACUAG
crD10MM9: CCUACGCCUUCAGCUCCAACUAG
crD10MM11: CCUACGCCAUGAGCUCCAACUAG
crD10MM12: CCUACGCCAUCUGCUCCAACUAG
crD10MM13: CCUACGCCAUCACCUCCAACUAG
crD10MM14: CCUACGCCAUCAGGUCCAACUAG
crD10MM15: CCUACGCCAUCAGCACCAACUAG
crD10MM16: CCUACGCCAUCAGCUGCAACUAG
crD10MM17: CCUACGCCAUCAGCUCGAACUAG
crD10MM18: CCUACGCCAUCAGCUCCUACUAG
crD10MM19: CCUACGCCAUCAGCUCCAUCUAG
crD10MM20: CCUACGCCAUCAGCUCCAAGUAG
crD10MM21: CCUACGCCAUCAGCUCCAACAAG
crD10MM22: CCUACGCCAUCAGCUCCAACUUG
crD10MM23: CCUACGCCAUCAGCUCCAACUAC
B
c.34 variants:
G>C (G12R)
G>A (G12S)
G>T (G12C)
c.35 variants:
G>C (G12A)
G>A (G12D)
G>T (G12V)
KRAS mRNA
n = 1
crD10C11 vs c.34
n = 1
crD10C11 vs c.35
n = 2
crD10C11 vs WT
C
crNT
crD10C11
crD10C11MM9
crD10C11MM9
KRAS^WT
KRAS^G12D
KRAS^G12C
D
crD10C11-MM12
KRAS^WT mRNA
3'
WT
5'
ACGGAUGCGGUGGUCGAGGUUGAUGGU
5' (pos. 1) - CCUACGCCA U A U GCUCCAACUAC - 3' (pos. 23)
crD10C11-MM12
n_mismatch = 3
KRAS^WT mRNA
3'
G12D
5'
ACGGAUGCGGUAGUCGAGGUUGAUGGU
5' (pos. 1) - CCUACGCCAU AU GCUCCAACUAC - 3' (pos. 23)
crD10C11-MM12
n_mismatch = 2
KRAS^WT mRNA
3'
G12C
5'
ACGGAUGCGGUGUUCGAGGUUGAUGGU
5' (pos. 1) - CCUACGCCA U A U GCUCCAACUAC - 3' (pos. 23)
crD10C11-MM12
n_mismatch = 2

**Figure S1 | Supplementary data related to RfxCas13d-mediated silencing of KRAS G12C and G12D. (A)** Barplot showing silencing efficiency of single-nucleotide mismatched spacers targeting KRAS G12D against KRAS-WT (grey) vs KRAS G12D (dark blue) or KRAS G12C (light blue bars; lower), normalised against crNT at 48h post-transfection (at least n=3 independent experiments). Sequences for each crRNA are shown below the barplot: the "U" nucleotide (in dark blue) indicates the uracil in the spacer sequence that basepairs with the G12D and the orange nucleotides indicate the mismatched spacer nucleotide. **(B)** Schematic depicting bi- specific G12-targeting crRNAs. Due to sequence complementarity at the c.34 and c.35 positions of the KRAS transcript, these bi-specific crRNAs will always have at least a one-nt mismatch with any other G12 variant, but at least two mismatches with KRAS wildtype. **(C)** Representative fluorescence micrographs (10x magnification) showing silencing efficiency of $crC_{10}D_{11}$-$MM_{9}$ and $crC_{10}D_{11}$-$MM_{12}$, relative to non-selective crC10D11, at 48h post-transfection (companion data to Figure 1D). **(D)** Schematic depicting the sequence and base-pairing configuration of SNV-selective $crC_{10}D_{11}$-$MM_{12}$ with WT vs G12D- and G12C-mutant KRAS mRNA targets. All experiments performed in HEK293T overexpression models.

#### Figure S2

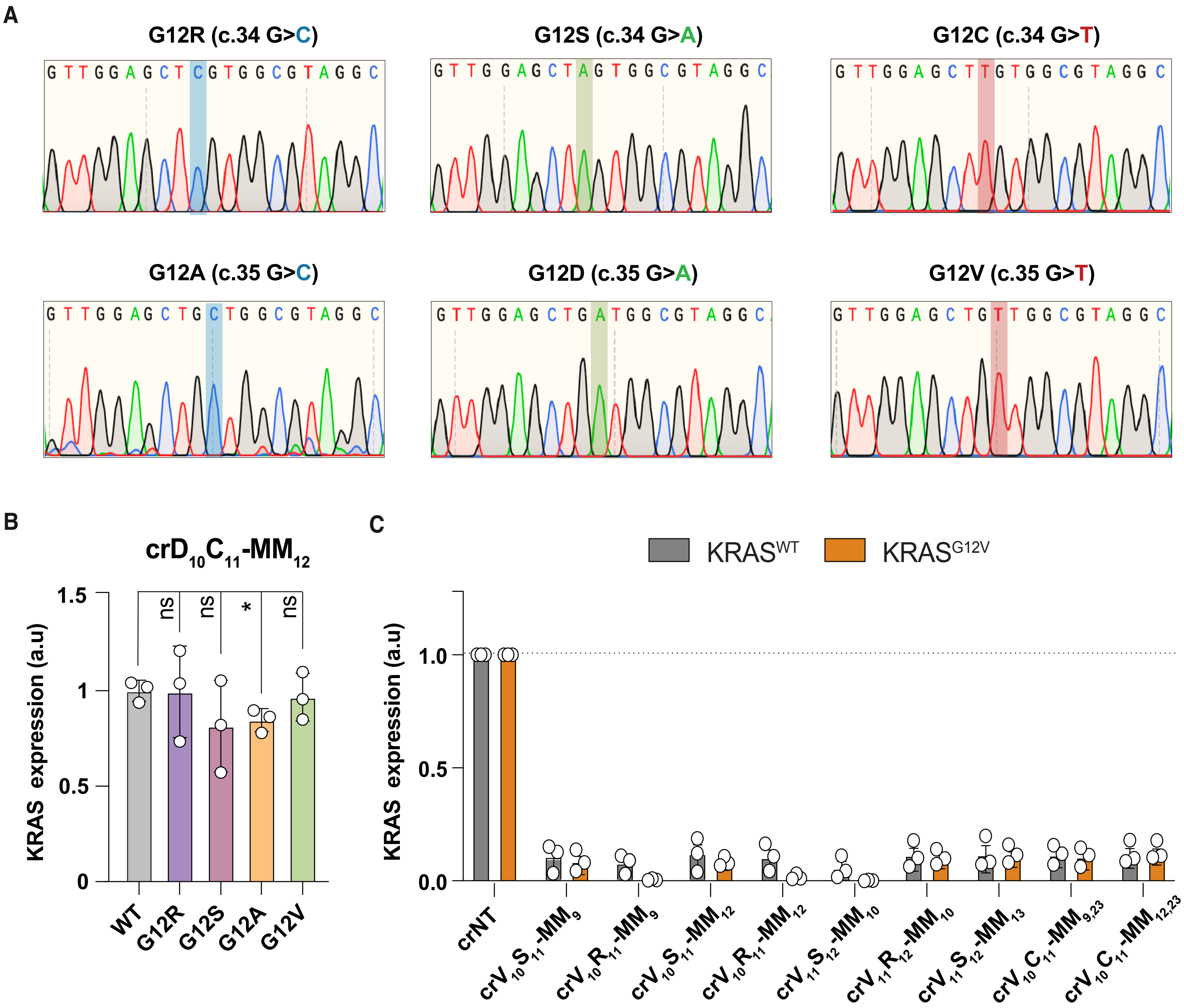


**Figure S2 | Supplementary data related to RfxCas13d-mediated silencing of additional KRAS G12 variants.** **(A)** Chromatograms from Sanger sequencing of the G12 hotspot plasmid library produced via site-directed mutagenesis of a KRAS-WT template, with knock-in SNVs highlighted. **(B)** Barplot showing minimal cross-reactive silencing by crC$_{10}$D$_{11}$-MM$_{12}$ against other c.34 and c.35 KRAS variants (G12R, purple; G12S, pink; G12A, green; and G12V, gold), normalised against the silencing of WT KRAS, at 48h post-transfection. **(C)** Barplot showing non-selective silencing by a panel of crC$_{10}$D$_{11}$-MM$_{9}$ and crC$_{10}$D$_{11}$-MM$_{12}$ derivatives against KRAS G12V (orange) vs KRAS WT (grey), normalised against crNT, at 48h post-transfection. All experiments performed in HEK293T overexpression models. For all bargraphs, individual points show the averaged mean fluorescence intensity from four fields-of-view per well, and error bars represent mean ± SD from three independent experiments. Statistical significance, where shown, was determined using unpaired t-tests, where * = $p < 0.05$, ** = $p < 0.01$, *** = $p < 0.001$, ****= $p < 0.0001$.

### Figure S3

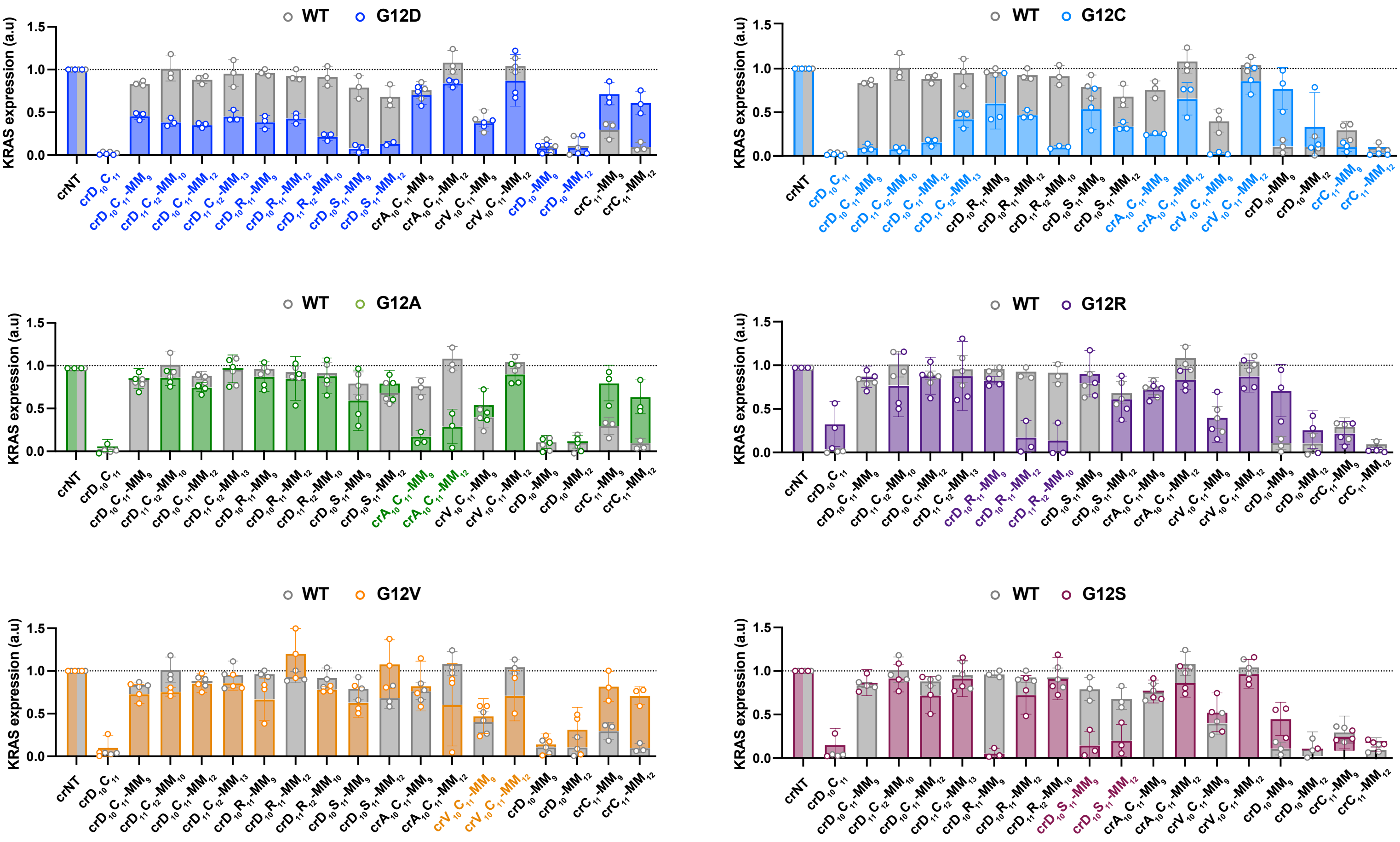


**Figure S3 | Silencing activity of extended crRNA mutagenesis library against all G12 hotspot mutations.** Barplots showing varying extents of cross-reactive silencing by $crC_{10}D_{11}$- $MM_9$ and $crC_{10}D_{11}$- $MM_{12}$ derivatives against all other c.34 and c.35 KRAS variants (G12D, dark blue; G12C, light blue; G12A, green; G12R, purple; G12V, gold; and G12S, pink), normalised against crNT, at 48h post-transfection. The coloured guides (on x-axis) indicate those which basepair with the corresponding variant and are therefore expected to show high(er) silencing activity. Data from n=3 independent experiments performed in HEK293T overexpression models. For all bargraphs, individual points show the averaged fluorescence intensity from 8 fields-of-view per experiment, and error bars represent mean ± SD from three independent experiments.

**Figure S4**

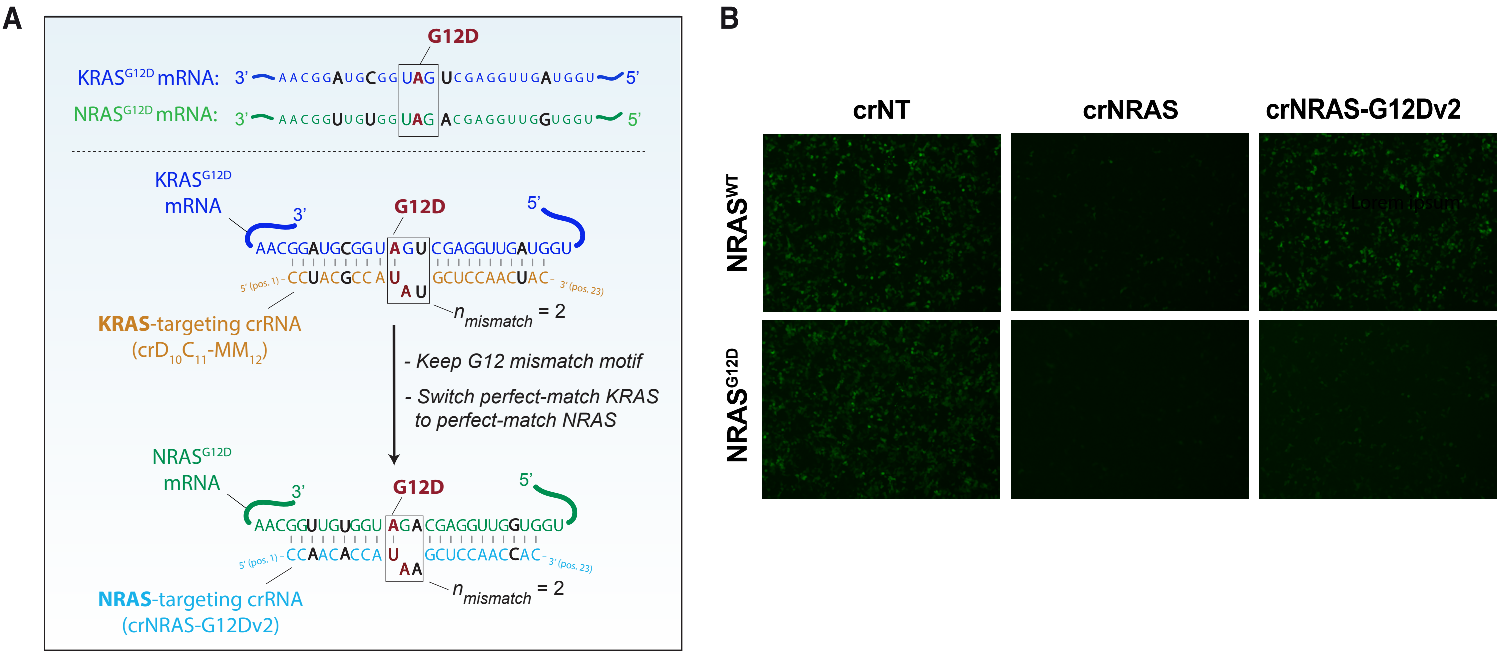


**Figure S2 | Supplementary data related to RfxCas13d-mediated silencing of NRAS G12D. (A)** Schematic of the guide design strategy used to adapt KRAS-G12D targeting crRNAs to NRAS-G12D targeting crRNAs, showing sequence homology between the two mRNAs (upper) the sequence of each crRNA (lower). The box indicates the nucleotides surrounding the codon 12 hotspot, and nucleotides in black indicate bases which are not conserved between KRAS and NRAS wildtype sequences. **(B)** Representative fluorescence micrographs (10x magnification) showing silencing efficiency of crNRAS-G12Dv2, relative to non-selective crNRAS and crNT control, at 48h post-transfection. All experiments were performed in HEK293T overexpression models.

**Figure S5**

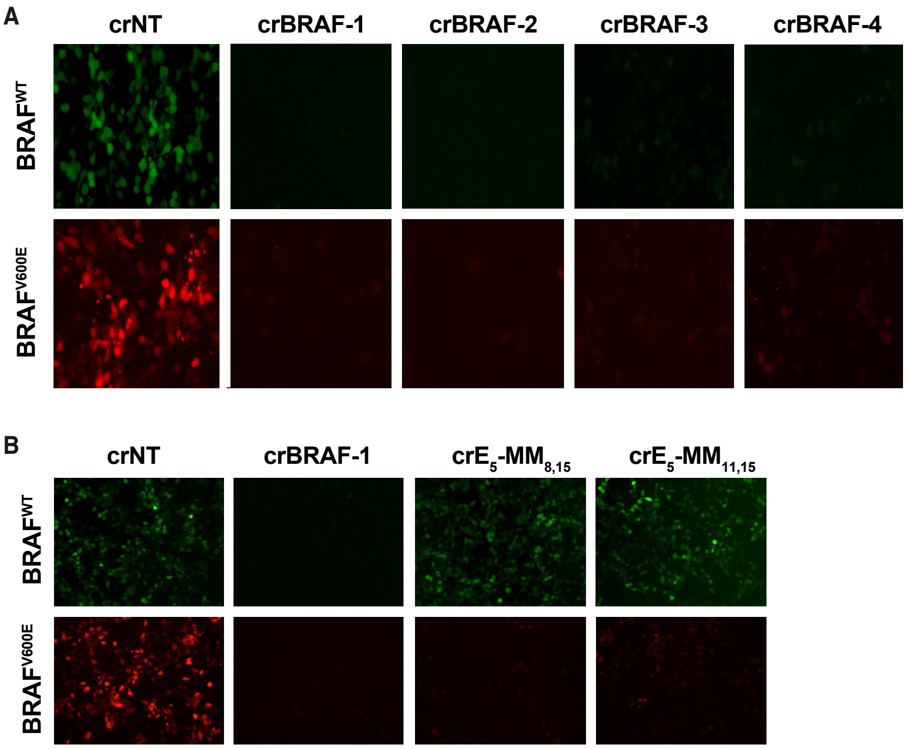


**Figure S5 | Supplementary data related to PspCas13b-mediated silencing of BRAF V600E. (A)** Representative fluorescence micrographs (20x magnification) showing silencing efficiency of four perfect-match crRNAs (crBRAF1-4) against BRAF WT vs V600E, normalised to a non- targeting control (crNT), at 48h post-transfection (companion data to Figure 4A). **(B)** Representative fluorescence micrographs (10x magnification) showing silencing efficiency of $crE_5$-$MM_{8,15}$ and $crE_5$-$MM_{11,15}$, relative to non-selective crBRAF-1, normalised to crNT control, at 48h post-transfection.

**Table S1: crRNA spacer sequences used in this study**

| **crRNA Name** | **CRISPR-Cas Orthologue** | **crRNA sequence** | **Target SNV** | **Figure** |
|---|---|---|---|---|
| crNT | RfxCas13d | UAGAUUGCUGUUCUACCAAGUGG | N/A | Multiple |
| crD10 | RfxCas13d | CCUACGCCAUCAGCUCCAACUAC | G12D | S1A |
| crD10-MM1 | RfxCas13d | GCUACGCCAUCAGCUCCAACUAC | G12D | S1A |
| crD10-MM2 | RfxCas13d | CGUACGCCAUCAGCUCCAACUAC | G12D | S1A |
| crD10-MM3 | RfxCas13d | CCAACGCCAUCAGCUCCAACUAC | G12D | S1A |
| crD10-MM4 | RfxCas13d | CCUUCGCCAUCAGCUCCAACUAC | G12D | S1A |
| crD10-MM5 | RfxCas13d | CCUAGGCCAUCAGCUCCAACUAC | G12D | S1A |
| crD10-MM6 | RfxCas13d | CCUACCCCAUCAGCUCCAACUAC | G12D | S1A |
| crD10-MM7 | RfxCas13d | CCUACGGCAUCAGCUCCAACUAC | G12D | S1A |
| crD10-MM8 | RfxCas13d | CCUACGCGAUCAGCUCCAACUAC | G12D | S1A |
| crD10-MM9 | RfxCas13d | CCUACGCCUUCAGCUCCAACUAC | G12D | S1A |
| crD10-MM11 | RfxCas13d | CCUACGCCAUGAGCUCCAACUAC | G12D | S1A |
| crD10-MM12 | RfxCas13d | CCUACGCCAUCUGCUCCAACUAC | G12D | S1A |
| crD10-MM13 | RfxCas13d | CCUACGCCAUCACCUCCAACUAC | G12D | S1A |
| crD10-MM14 | RfxCas13d | CCUACGCCAUCAGGUCCAACUAC | G12D | S1A |
| crD10-MM15 | RfxCas13d | CCUACGCCAUCAGCACCAACUAC | G12D | S1A |
| crD10-MM16 | RfxCas13d | CCUACGCCAUCAGCUGCAACUAC | G12D | S1A |
| crD10-MM17 | RfxCas13d | CCUACGCCAUCAGCUCGAACUAC | G12D | S1A |
| crD10-MM18 | RfxCas13d | CCUACGCCAUCAGCUCCUACUAC | G12D | S1A |
| crD10-MM19 | RfxCas13d | CCUACGCCAUCAGCUCCAUCUAC | G12D | S1A |
| crD10-MM20 | RfxCas13d | CCUACGCCAUCAGCUCCAAGUAC | G12D | S1A |
| crD10-MM21 | RfxCas13d | CCUACGCCAUCAGCUCCAACAAC | G12D | S1A |
| crD10-MM22 | RfxCas13d | CCUACGCCAUCAGCUCCAACUUC | G12D | S1A |
| crD10-MM23 | RfxCas13d | CCUACGCCAUCAGCUCCAACUAG | G12D | S1A |
| crD10C11 | RfxCas13d | CCUACGCCAUAAGCUCCAACUAC | G12C, G12D | Multiple |
| crC11-MM23 | RfxCas13d | CCUACGCCACAAGCUCCAACUAG | G12C | 1C, 2F |
| crD10C11-MM1 | RfxCas13d | GCUACGCCAUAAGCUCCAACUAC | G12C, G12D | 1C, 2F |
| crD10C11-MM4 | RfxCas13d | GCUUCGCCAUAAGCUCCAACUAC | G12C, G12D | 1C, 2F |
| crD10C11-MM6 | RfxCas13d | GCUACCCCAUAAGCUCCAACUAC | G12C, G12D | 1C, 2F |
| crD10C11-MM9 | RfxCas13d | GCUACGCCUUAAGCUCCAACUAC | G12C, G12D | Multiple |
| crD10C11-MM12 | RfxCas13d | CCUACGCCAUAUGCUCCAACUAC | G12C, G12D | Multiple |
| crD10C11-MM15 | RfxCas13d | CCUACGCCAUAAGCACCAACUAC | G12C, G12D | 1C, 2F |
| crD10C11-MM18 | RfxCas13d | CCUACGCCAUAAGCUCCUACUAC | G12C, G12D | 1C, 2F |
| crD10C11-MM21 | RfxCas13d | CCUACGCCAUAAGCUCCAACAAC | G12C, G12D | 1C, 2F |
| crD10C11-MM23 | RfxCas13d | CCUACGCCAUAAGCUCCAACUAG | G12C, G12D | 1C, 2F |
| crD10C11-MM1,23 | RfxCas13d | GCUACGCCAUAAGCUCCAACUAG | G12C, G12D | 1C, 2F |
| crD10C11-MM4,23 | RfxCas13d | CCUUCGCCAUAAGCUCCAACUAG | G12C, G12D | 1C, 2F |
| crD10C11-MM6,23 | RfxCas13d | CCUACCCCAUAAGCUCCAACUAG | G12C, G12D | 1C, 2F |
| crD10C11-MM9,23 | RfxCas13d | CCUACGCCUUAAGCUCCAACUAG | G12C, G12D | 1C, 2F |
| crD10C11-MM12,23 | RfxCas13d | CCUACGCCAUAUGCUCCAACUAG | G12C, G12D | 1C, 2F |
| crD10C11-MM15,23 | RfxCas13d | CCUACGCCAUAAGCACCAACUAG | G12C, G12D | 1C, 2F |
| crD10C11-MM18,23 | RfxCas13d | CCUACGCCAUAAGCUCCUACUAG | G12C, G12D | 1C, 2F |
| crD10C11-MM21,23 | RfxCas13d | CCUACGCCAUAAGCUCCAACAAG | G12C, G12D | 1C, 2F |
| crD11C12-MM10 **[Alias: crG12C]** | RfxCas13d | GCCUACGCCUUAAGCUCCAACUA | G12C, G12D | Multiple |
| crD11C12-MM13 | RfxCas13d | GCCUACGCCAUAUGCUCCAACUA | G12C, G12D | 2E, 2F |
| crC11-MM9 | RfxCas13d | CCUACGCCUCAAGCUCCAACUAC | WT, G12C | 2F |
| crC11-MM12 | RfxCas13d | CCUACGCCACAUGCUCCAACUAC | WT, G12C | 2F |
| crA10C11-MM9 **[Alias: crG12A]** | RfxCas13d | CCUACGCCUGAAGCUCCAACUAC | G12C, G12A | Multiple |
| crA10C11-MM12 | RfxCas13d | CCUACGCCAGAUGCUCCAACUAC | G12C, G12A | 2D-2F |
| crV10C11-MM12 | RfxCas13d | CCUACGCCAAAUGCUCCAACUAC | G12C,G12V | 2D-2F |
| crV10C11-MM9 | RfxCas13d | CCUACGCCUAAAGCUCCAACUAC | G12C,G12V | 2D-2F |
| crV10R11-MM9 | RfxCas13d | CCUACGCCUAGAGCUCCAACUAC | G12R, G12V | S2C, 2F |
| crV10R11-MM12 | RfxCas13d | CCUACGCCAAGUGCUCCAACUAC | G12R, G12V | S2C, 2F |
| crV10S11-MM9 | RfxCas13d | CCUACGCCUAUAGCUCCAACUAC | G12S, G12V | S2C, 2F |
| crV10S11-MM12 | RfxCas13d | CCUACGCCAAUUGCUCCAACUAC | G12S, G12V | 2D-2F |
| crD11-R12-MM10 **[Alias: crG12R]** | RfxCas13d | GCCUACGCCUUGAGCUCCAACUA | G12D, G12R | Multiple |
| crD11-MM9,10 | RfxCas13d | GCCUACGCGUUCAGCUCCAACUA | G12D | 2D-2F |
| crV11S12-MM10 | RfxCas13d | GCCUACGCCUAUAGCUCCAACUA | G12S, G12V | S2C, 2F |
| crV11R12-MM10 | RfxCas13d | GCCUACGCCUAGAGCUCCAACUA | G12R, G12V | S2C, 2F |
| crV11S12-MM13 | RfxCas13d | GCCUACGCCAAUUGCUCCAACUA | G12S, G12V | S2C, 2F |
| crV10C11-MM9,23 | RfxCas13d | CCUACGCCUAAAGCUCCAACUAG | G12C, G12V | S2C, 2F |
| crV10C11-MM12,23 | RfxCas13d | CCUACGCCAAAUGCUCCAACUAG | G12C, G12V | S2C, 2F |
| crD10R11-MM12 | RfxCas13d | CCUACGCCAUGUGCUCCAACUAC | G12D, G12R | 2D-2F |
| crD10R11-MM9 | RfxCas13d | CCUACGCCUUGAGCUCCAACUAC | G12D, G12R | 2D-2F |
| crD10S11-MM12 | RfxCas13d | CCUACGCCAUUUGCUCCAACUAC | G12D, G12S | 2D-2F |
| crD10S11-MM9 **[Alias: crG12D & crD12S]** | RfxCas13d | CCUACGCCUUUAGCUCCAACUAC | G12D, G12S | Multiple |
| crNRAS | RfxCas13d | CCAACACCATCTGCTCCAACCAC | NRAS WT, NRAS G12D | 3F, 3G |
| crNRAS-G12Dv1 | RfxCas13d | CCAACACCUUAUGCUCCAACCAC | NRAS G12D | 3F, 3G |
| crNRAS-G12Dv2 | RfxCas13d | CCAACACCAUAAGCUCCAACCAC | NRAS G12D | 3F, 3G |
| crNT | PspCas13b | UAGAUUGCUGUUCUACCAAGUAAUCCAUCA | V600E | Multiple |
| crBRAF-1 | PspCas13b | GGUCUCUGUAGCUAGACCAAAAUCACCUAU | V600E | Multiple |
| crBRAF-2 | PspCas13b | GGGAGAUUUCUCUGUAGCUAGACCAAAAUCA | V600E | 4A, S4A |
| crBRAF-3 | PspCas13b | GGUCCAUCGAGAUUUCUCUGUAGCUAGACC | V600E | 4A, S4A |
| crBRAF-4 | PspCas13b | GGACCCACUCCAUCGAGAUUUCUCUGUAGC | V600E | 4A, S4A |
| crE5-MM1-2(v1) | PspCas13b | GGAGUCUGUAGCUAGACCAAAAUCACCUAU | V600E | 4B |
| crE5-MM1-2(v2) | PspCas13b | GGCGUCUGUAGCUAGACCAAAAUCACCUAU | V600E | 4B |
| crE5-MM6-7(v1) | PspCas13b | GGUCUGAGUAGCUAGACCAAAAUCACCUAU | V600E | 4B |
| crE5-MM6-8(v1) | PspCas13b | GGUCUGACUAGCUAGACCAAAAUCACCUAU | V600E | 4B |
| crE5-MM6-9(v1) | PspCas13b | GGUCUGACAAGCUAGACCAAAAUCACCUAU | V600E | 4B |
| crE5-MM6-7(v2) | PspCas13b | GGUCUGCGUAGCUAGACCAAAAUCACCUAU | V600E | 4B |
| crE5-MM6-8(v2) | PspCas13b | GGUCUGCCUAGCUAGACCAAAAUCACCUAU | V600E | 4B |
| crE5-MM6-9(v2) | PspCas13b | GGUCUGCCCAGCUAGACCAAAAUCACCUAU | V600E | 4B |
| crE5-MM4,8 | PspCas13b | GGUGUCUCUAGCUAGACCAAAAUCACCUAU | V600E | 4B |
| crE5-MM4,11 | PspCas13b | GGUGUCUGUACCUAGACCAAAAUCACCUAU | V600E | 4B |
| crE5-MM4,15 | PspCas13b | GGUGUCUGUAGCUACACCAAAAUCACCUAU | V600E | 4B |
| crE5-MM8,11 | PspCas13b | GGUCUCUCUACCUAGACCAAAAUCACCUAU | V600E | 4B |
| crE5-MM8,15 | PspCas13b | GGUCUCUCUAGCUACACCAAAAUCACCUAU | V600E | Multiple |
| crE5-MM11,15 | PspCas13b | GGUCUCUGUACCUACACCAAAAUCACCUAU | V600E | Multiple |
| crE5-MM8,11,15 | PspCas13b | GGUCUCUCUACCUACACCAAAAUCACCUAU | V600E | 4B |
| crE5-MM4,15,26 | PspCas13b | GGUGUCUGUAGCUACACCAAAAUCAGCUAU | V600E | 4B |
| crE5-MM8,15,22 | PspCas13b | GGUCUCUCUAGCUACACCAAAUUCACCUAU | V600E | 4B |
| crE5-MM4,11,18,25 | PspCas13b | GGUGUCUGUACCUAGACGAAAAUCUCCUAU | V600E | 4B |
| crE5-MM6,13,20,26 | PspCas13b | GGUCUGUGUAGCAAGACCAUAAUCAGCUAU | V600E | 4B |
| crE5-MM8,15,22,28 | PspCas13b | GGUCUCUGUAGCUACACCAAAUUCACCAAU | V600E | 4B |

**Table S2: Primers used for Sanger sequencing, site-directed mutagenesis an[illegible]**

| Primer Name (Forward / Reverse) | Sequence |
|---|---|
| IRES (R) | GCATTCCTTTGGCGAGAG |
| M13 (R) | CAGGAAACAGCTATGAC |
| GAPDH (F) | GGAGCGAGATCCCTCCAAAAT |
| GAPDH (R) | GGCTGTTGTCA T ACTTCTCA TGG |
| BRAF (F) | CCATATCATTGAGACCAAATTTGAGATG |
| BRAF (R) | GGCACTCTGCCATTAATCTCTTCATGG |
| KRAS G12A (F) | TGTGGTAGTTGGAGCTGCTGGCGTAGGCAAGAGTG |
| KRAS G12A (R) | CACTCTTGCCTACGCCAGCAGCTCCAACTACCACA |
| KRAS G12C (F) | TTGTGGTAGTTGGAGCTTGTGGCGTAGGCAAGAGT |
| KRAS G12C (R) | ACTCTTGCCTACGCCACAAGCTCCAACTACCACAA |
| KRAS G12D (F) | TGTGGTAGTTGGAGCTGATGGCGTAGGCAAGAGTG |
| KRAS G12D (R) | CACTCTTGCCTACGCCATCAGCTCCAACTACCACA |
| KRAS G12R (F) | TTGTGGTAGTTGGAGCTCGTGGCGTAGGCAAGAGT |
| KRAS G12R (R) | ACTCTTGCCTACGCCACGAGCTCCAACTACCACAA |
| KRAS G12S (F) | TTGTGGTAGTTGGAGCTAGTGGCGTAGGCAAGAGT |
| KRAS G12S (R) | ACTCTTGCCTACGCCACTAGCTCCAACTACCACAA |
| KRAS G12V (F) | TGTGGTAGTTGGAGCTGTTGGCGTAGGCAAGAGTG |
| KRAS G12V (R) | CACTCTTGCCTACGCCAACAGCTCCAACTACCACA |

**Table S3:Antibodies used for WB**

| Target (Clone) | Type | Host Species | Reactivity |
|---|---|---|---|
| Anti-HA (6E2) | Primary | Mouse | N/A (species agnostic) |
| Anti-FLAG (M2) | Primary | Mouse | N/A (species agnostic) |
| Anti-tubulin (polyclonal) | Primary | Rabbit | Human |
| Anti-mouse IgG | Secondary | Goat | Mouse |
| Anti-rabbit IgG | Secondary | Donkey | Rabbit |
